# Quantifying transmission of emerging zoonoses: Using mathematical models to maximize the value of surveillance data

**DOI:** 10.1101/677021

**Authors:** Monique R. Ambrose, Adam J. Kucharski, Pierre Formenty, Jean-Jacques Muyembe-Tamfum, Anne W. Rimoin, James O. Lloyd-Smith

**Author notes:** Corresponding author (JOL-S).

## Abstract

Understanding and quantifying the transmission of zoonotic pathogens is essential for directing public health responses, especially for pathogens capable of transmission between humans. However, determining a pathogen’s transmission dynamics is complicated by challenges often encountered in zoonotic disease surveillance, including unobserved sources of transmission (both human and zoonotic), limited spatial information, and unknown scope of surveillance. In this work, we present a model-based inference method that addresses these challenges for subcritical zoonotic pathogens using a spatial model with two levels of mixing. After demonstrating the robustness of the method using simulation studies, we apply the new method to a dataset of human monkeypox cases detected during an active surveillance program from 1982-1986 in the Democratic Republic of the Congo (DRC). Our results provide estimates of the reproductive number and spillover rate of monkeypox during this surveillance period and suggest that most human-to-human transmission events occur over distances of 30km or less. Taking advantage of contact-tracing data available for a subset of monkeypox cases, we find that around 80% of contact-traced links could be correctly recovered from transmission trees inferred using only date and location. Our results highlight the importance of identifying the appropriate spatial scale of transmission, and show how even imperfect spatiotemporal data can be incorporated into models to obtain reliable estimates of human-to-human transmission patterns.

**Author Summary:** Surveillance datasets are often the only sources of information about the ecology and epidemiology of zoonotic infectious diseases. Methods that can extract as much information as possible from these datasets therefore provide a key advantage for informing our understanding of the disease dynamics and improving our ability to choose the optimal intervention strategy. We developed and tested a likelihood-based inference method based on a mechanistic model of the spillover and human-to-human transmission processes. We first used simulated datasets to explore which information about the disease dynamics of a subcritical zoonotic pathogen could be successfully extracted from a line-list surveillance dataset with non-localized spatial information and unknown geographic coverage. We then applied the method to a dataset of human monkeypox cases detected during an active surveillance program in the Democratic Republic of the Congo between 1982 and 1986 to obtain estimates of the reproductive number, spillover rate, and spatial dispersal of monkeypox in humans.

## Introduction

Many recent infectious disease threats have been caused by pathogens with zoonotic origins, including Ebola, pandemic H1N1 influenza, and SARS- and MERS-Coronaviruses, and zoonotic pathogens are expected to be a primary source of future emerging infectious diseases [1–8]. By definition, zoonotic pathogens can transmit from animals to humans; those also capable of human-to-human transmission are of particular public health concern [5,9]. Infectious disease surveillance serves a crucial role for detecting and gathering information on zoonotic pathogens: data obtained through surveillance are often the primary resource available for informing public health management decisions [10]. Developing methods that improve our ability to infer information about a pathogen’s transmission dynamics from available surveillance data is therefore an essential frontier for understanding and ultimately combating these pathogens [11,12].

For zoonoses, three epidemiological measures are crucial for summarizing transmission dynamics and informing risk assessments. The first of these is the spillover rate, which indicates how frequently the pathogen is transmitted from the animal reservoir into humans and helps inform the total expected disease incidence [13]. The second measure describes the pathogen’s potential for further spread once in the human population and is commonly assessed using the reproductive number (*R*), which gives the average number of secondary human cases caused by an infectious individual [14,15]. Values of *R* greater than one indicate that the pathogen is capable of sustained (i.e. ‘supercritical’) transmission in humans. Pathogens with subcritical transmission (*R* less than one but greater than zero) can cause limited chains of transmission in humans after a zoonotic introduction, and they pose a risk of acquiring ability for supercritical transmission via evolutionary or environmental change [2,5,16]. The third epidemiological measure is the distance over which human-to-human transmission occurs, which informs how the disease will spread spatially and the risk of it being introduced into new populations. Combined, these three measures can help evaluate the current public health threat posed by the pathogen, the risk of future emergence, and the most effective approaches for disease management.

Estimating epidemiological measures is a challenging task in any pathogen system, and the unique properties of zoonotic diseases can exacerbate these difficulties. Infectious disease surveillance often records temporal information and certain aspects of spatial information about human cases, but the underlying transmission events are seldom observed. In a zoonotic system, this means that an observed human infection could have been caused by a previous human case or by zoonotic spillover. Without intensive contact tracing, or sequence data in the case of fast-evolving pathogens, quantifying the relative contribution of zoonotic versus human-to-human transmission is a major challenge; identifying the source of infection for specific individuals is an even bigger one.

Epidemiological analyses are often hindered by data truncation and unknown denominators [17,18]. In many disease surveillance systems, the total set of localities under surveillance (i.e. those that would appear in the dataset if a case occurred there) can be separated into ‘observed localities,’ which appear in the dataset because they reported one or more cases, and ‘silent localities,’ which have no cases during the surveillance period and therefore do not appear in the dataset. This form of truncation, where localities with zero cases are absent from the dataset, obscures the true scope of the surveillance effort. Without knowledge of the total number of localities under observation (the ‘unknown denominator’), accurately estimating the spillover rate and probability of human-to-human transmission between localities is not straightforward. Simply disregarding these silent localities in the analysis is the functional equivalent of selectively removing zeros from the dataset and can lead to problematic inference biases.

Complicating inference efforts further is the fact that surveillance datasets often report the geographic location of cases only at a coarse resolution, obscuring information about a transmission process that occurs on a much finer scale [19–21]. Precise spatial information is often absent from historic datasets and data collected in remote or low-resource areas, replaced by the names of the locality and broader administrative units where the case occurred. For example, only the village name and the region and country to which the village belongs may be recorded in a dataset. Furthermore, linking a village name to spatial coordinates is often impossible when maps of the region do not exist or only unofficial local names are used. Although collecting exact spatial coordinates has become more practical in contemporary disease surveillance, privacy and confidentiality concerns can arise in both human and agricultural contexts when data contains high-resolution spatial information [19,20,22–25], leading to data being reported in a non-localized manner. Methods that can use this inexact spatial information are especially needed for zoonotic diseases, where any additional information about the proximity of human cases to one another can improve the power to distinguish between human-to-human transmission and zoonotic spillover.

Despite these challenges, a series of research efforts have expanded our ability to estimate the transmission properties of zoonotic pathogens from case onset data. A key set of methods revolve around inferring *R* from the sizes of case clusters (a cluster is defined as a group of cases that occur in close spatiotemporal proximity to one another) or from the proportion of observed cases that were infected by zoonotic spillover [16,26–30]. However, these approaches either require detailed case investigations to determine whether a case was infected by a zoonotic or human source or assume that each cluster is caused by one single spillover event followed by human-to-human transmission. A likelihood-based approach for estimating *R* for human-to-human transmission using only symptom onset dates of cases was introduced by Wallinga and Teunis [31]. This method was extended to apply to zoonotic systems by Lo Iacono et al. [32], but the extension requires that chains of exclusively human-to-human transmission can be identified, and is thus not applicable to many zoonotic surveillance systems where human and zoonotic transmissions are intermixed. A different approach was taken by White and Pagano [33], who introduced a different likelihood-based method that compares the observed number of cases on each day with the expected number, as calculated using the number and timing of previous cases. Though the White and Pagano approach was only applicable to human-to-human transmission, it was expanded by Kucharski et al. [34] to work in zoonotic spillover systems in scenarios where a control measure, implemented at a known point in time, causes an abrupt reduction in spillover. A related approach that requires knowledge of the human and animal reservoir population sizes was also explored in Lo Iacono et al. [35]. Crucially, however, none of these methods incorporate information about the spatial location of cases to improve inference power or to estimate patterns of spatial spread. Spatial data is a powerful tool in transmission inference in single-species studies (e.g. [36–39]), but has largely been excluded from analyses of zoonotic transmission, which often implicitly assume homogenous mixing across the study area or that human-to-human transmission can only occur within a locality. One recent exception to this is the analysis by Cauchemez et al. [40], which includes transmission at several spatial levels.

In this work, we present model-based inference methods that allow us to infer *R*, the spillover rate, and properties of spatial spread among humans from surveillance datasets with non-localized spatial information and an unknown total number of surveilled localities. Our approach builds on methods introduced by White and Pagano [33] and Kucharski et al. [34], but allows continuous spillover throughout the surveillance period and makes use of available spatial information on case location. While the method could be readily adjusted to incorporate more precise geographic information should it be available, in this study we focus on the more challenging scenario in which only the names of the locality and broader administrative units where a case occurred are known. To make use of this form of non-localized spatial data, our model considers two scales of spatial mixing and transmission (Fig 1A), reminiscent of the ‘epidemics with two levels of mixing’ structure utilized in Ball et al. [41] and Demiris and O’Neill [42]. The first mixing level is the locality in which the case occurred, such as a village, conceptualized as a group of individuals geographically separated from other localities. We assume that individuals within the same locality have more frequent contact with one another than with individuals from other localities, and therefore that infection is more likely to be transmitted within a locality. However, the total number of localities under surveillance is unknown because only localities with one or more cases appear in the dataset (the ‘unknown denominator’ problem discussed above). We refer to the second spatial level as the ‘broader contact zone.’ It describes a collection of localities that all occur within the same administrative unit and likely share some amount of human movement. When multiple types of administrative units of different sizes are reported in the dataset (e.g., districts, regions, provinces, etc.), the ideal choice for broader contact zone is the smallest administrative unit that contains inter-locality human-to-human transmission events. If this scale is not known *a priori*, inferring the appropriate scale of administrative unit is necessary.

**Fig 1.**
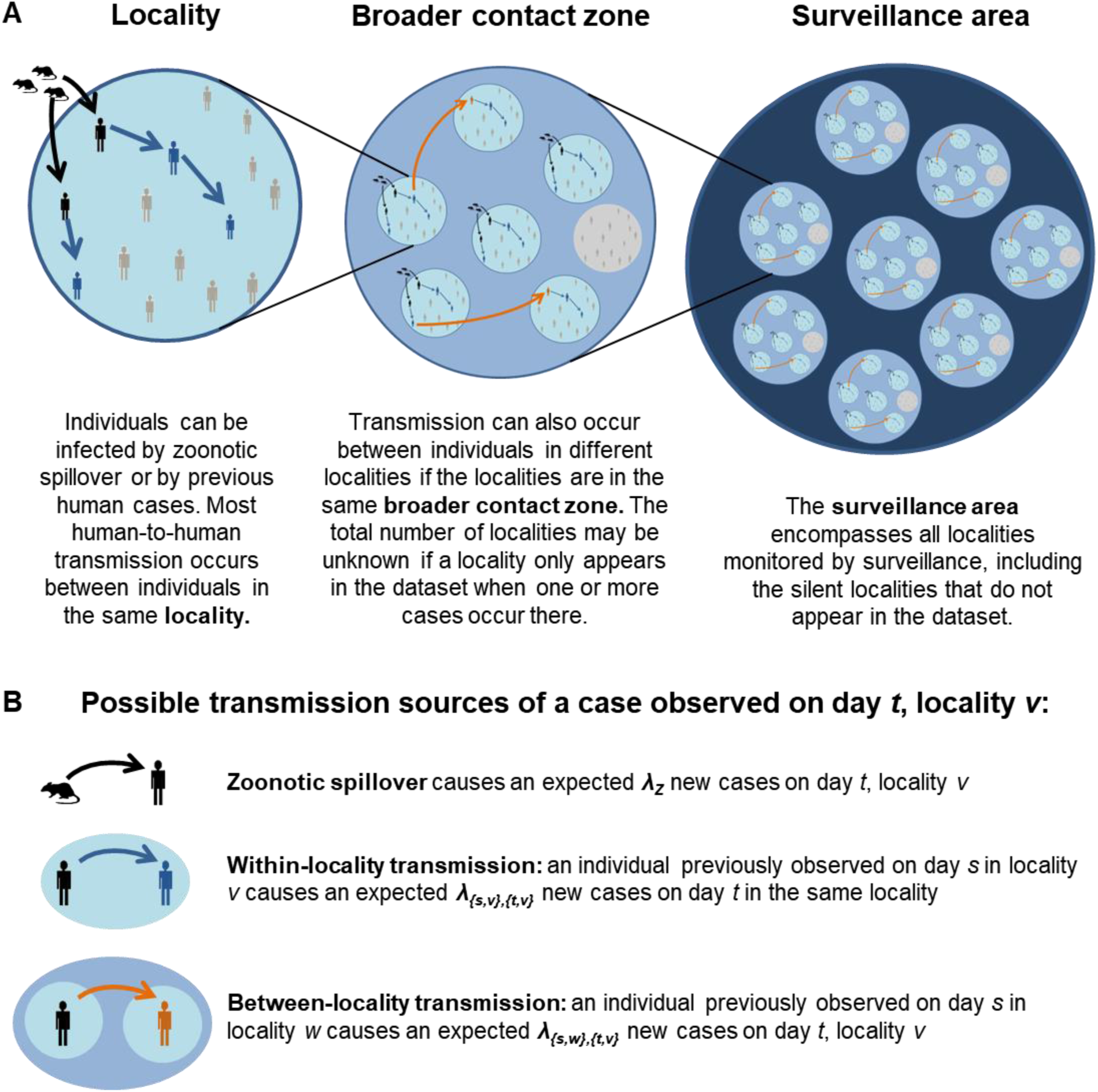
Model schematic. **A.** The schematic illustrates the spatial scales considered in the model and the types of transmission that occurs at different scales. Human cases are represented in black if they were infected by zoonotic spillover, blue if they were infected by within-locality human-to-human transmission, and orange if infected by between-locality human-to-human transmission. Individuals who are not infected are colored gray and do not appear in the surveillance dataset. Similarly, if zero individuals in a locality are infected, that ‘silent locality’ does not appear in the dataset (represented by the gray locality in the broader contact zone). **B.** The possible sources of human infection, which in aggregate determine the number of new infections on day *t*, locality *v*. The number of cases arising from spillover and human-to-human transmissions follow Poisson distributions with means *λ_Z_* and *λ_{s,w},{t,v}_*, respectively.

We tested the method against a variety of datasets simulated using different epidemiological parameters, offspring distributions for human-to-human transmission, and spatial transmission kernels. To assess the performance of the method, we compared the estimated and true values for epidemiological measures such as the reproductive number and spillover rate, and also examined how well the method was able to estimate the probable transmission source of each case. When silent localities were not accounted for, substantial biases arose in zoonotic spillover rate estimates. However, a modified method that accounts for these silent localities was successful in a wide range of circumstances. We therefore applied this ‘corrected-denominator method’ to a dataset on human monkeypox cases from an active surveillance effort conducted in the Democratic Republic of the Congo (formerly Zaire) in the 1980s [43] (Fig 2). Gaining insights to the disease dynamics of human monkeypox is particularly relevant given the recent increase in monkeypox incidence and outbreaks and the growing list of countries and regions reporting human monkeypox cases [44–51]. Using the high-coverage 1980s surveillance dataset to quantify the pathogen’s transmission dynamics will improve our understanding of what drives its spread and lays the groundwork to assess what has changed over the past decades to give rise to observed increases. With the 1980s monkeypox surveillance dataset, we repeated the analyses using four different assumptions about the appropriate spatial scale to represent the ‘broader contact zone’ over which human-to-human transmissions take place and selected the preferred option using the deviance information criterion (DIC) method for model comparison. In the monkeypox dataset, contact-tracing data are available for a subset of the cases, providing a rare opportunity to compare inferred transmission sources with those suggested by epidemiological investigation. In addition, some localities were associated with known GPS coordinates, enabling us to estimate the spatial transmission kernel in greater detail. As such, our monkeypox analysis yielded estimates of *R* and the spillover rate during the 1980s surveillance period, as well as insights into the spatial scale of human transmission of monkeypox.

**Fig 2.**
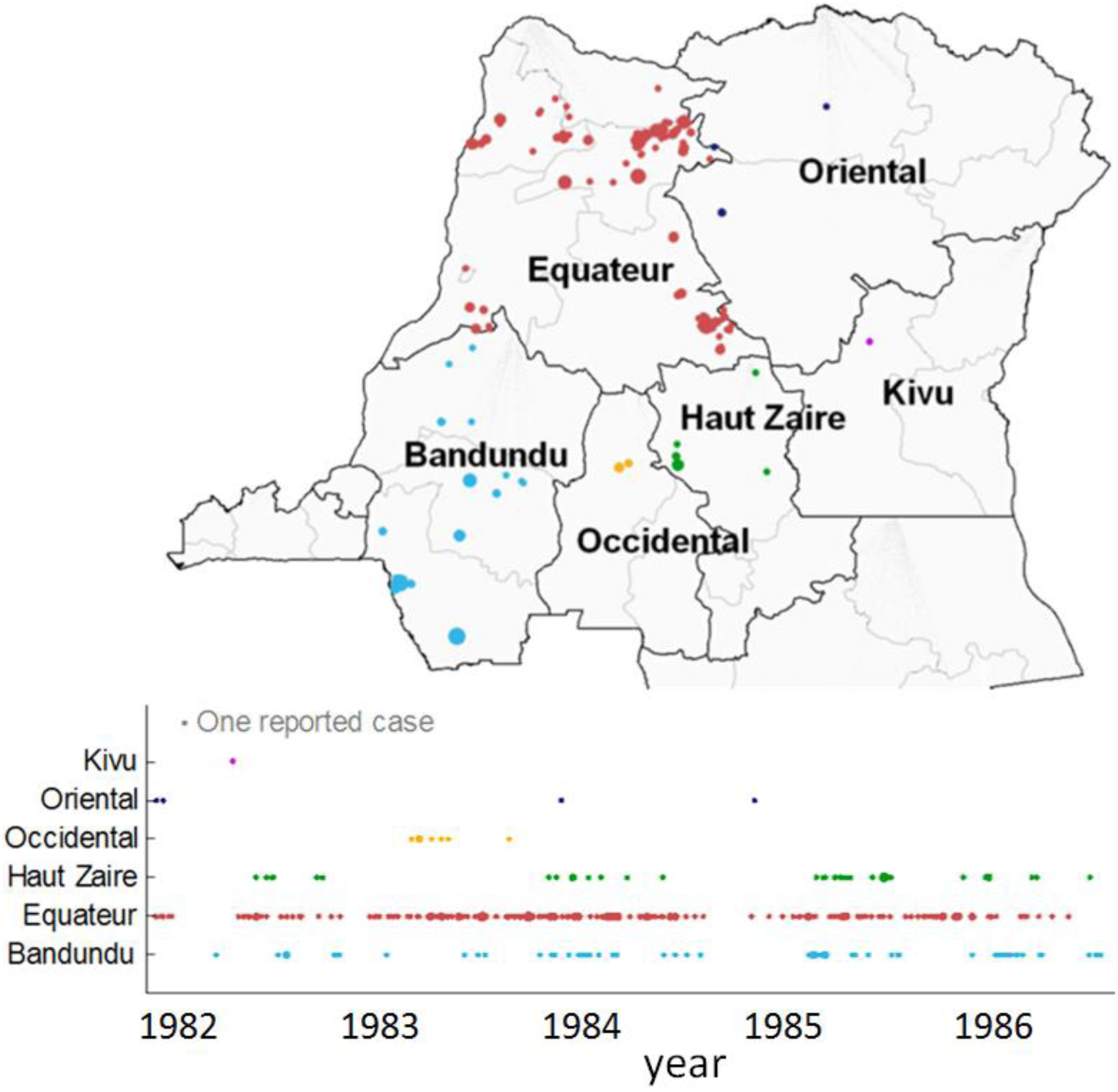
Map and time-series showing locations and dates of human monkeypox cases. The size of points on the map indicate the number of cases and the color of points corresponds to the region in which the cases occurred. Dark lines indicate region boundaries while light lines indicate the official boundaries for districts (though in the monkeypox surveillance dataset these are sometimes further divided into administrative subregions).

## Results

### Overview of the approach

We first validated the inference framework using a simulation study, then applied the validated method to a dataset on human monkeypox cases to estimate key epidemiological parameters and the spatial scale of transmission. To generate simulated test datasets and perform parameter inference, we used a mathematical model of the zoonotic pathogen’s transmission into and among humans. The model tracks the number of human cases that occur in each locality on each day; infections can arise from spillover from the zoonotic reservoir or from human-to-human transmission (Fig 1B). Three key parameters govern the behavior of the system. The spillover rate (*λ_z_*) describes the average number of human cases caused by animal-to-human transmission (‘primary cases’) in each locality per day. The reproductive number of the pathogen (*R*) determines the average number of (‘secondary’) cases caused by each infected human. And the spatial dispersal of the pathogen is controlled by the fraction of cases arising from human-to-human transmission that occur in the same locality as the source case (*σ*) and the rules governing inter-locality transmission events. Two spatial scales of transmission are included in the model: within the locality of the case and between localities in the same broader contact zone. Using this model (described further in Methods 4.1) and values for the three parameters, the likelihood of observing *N_t,v_* cases on each day *t* and locality *v* can be calculated. Markov chain Monte Carlo (MCMC) methods were used to infer posterior parameter distributions for a given dataset of cases.

### Robustness of model-based inference method

#### Basic method (assumes the total number of localities under surveillance is known)

To assess the accuracy and precision of our method’s estimates of spillover and transmission parameters, we simulated datasets with known parameter values and compared these true values with the inferred values. We investigated a range of *R* and *λ_z_* values in the neighborhood of values previously estimated for monkeypox [16,52], with *R* ranging from 0.2 to 0.6 and *λ_z_* ranging from 0.0001 to 0.0007 expected spillover events per locality per day (*λ_z_* values correspond to 59 to 415 expected spillover events in the five year simulation period, across all localities). Transmission events between humans had a probability *σ*=0.75 of occurring within a locality and otherwise were equally likely between any localities in the same broader contact zone. We were interested in seeing how well the inference methods are able to use the spatial-temporal arrangement of cases to estimate the true parameter values.

Across 125 simulations (25 simulations for each of five parameter sets), estimated values clustered around the true parameter values. The true value for *R* was included in the 95% credible interval (CI) 119 times (95.2%) and for *λ_z_* was included 121 times (96.8%) (Fig 3A). On average, the posterior mean estimate of *R* differed from the true value by 8.6%; the analogous percent errors for *λ_z_* and σ estimates were 6.3% and 7.0%, respectively (S1 Table).

**Fig 3.**
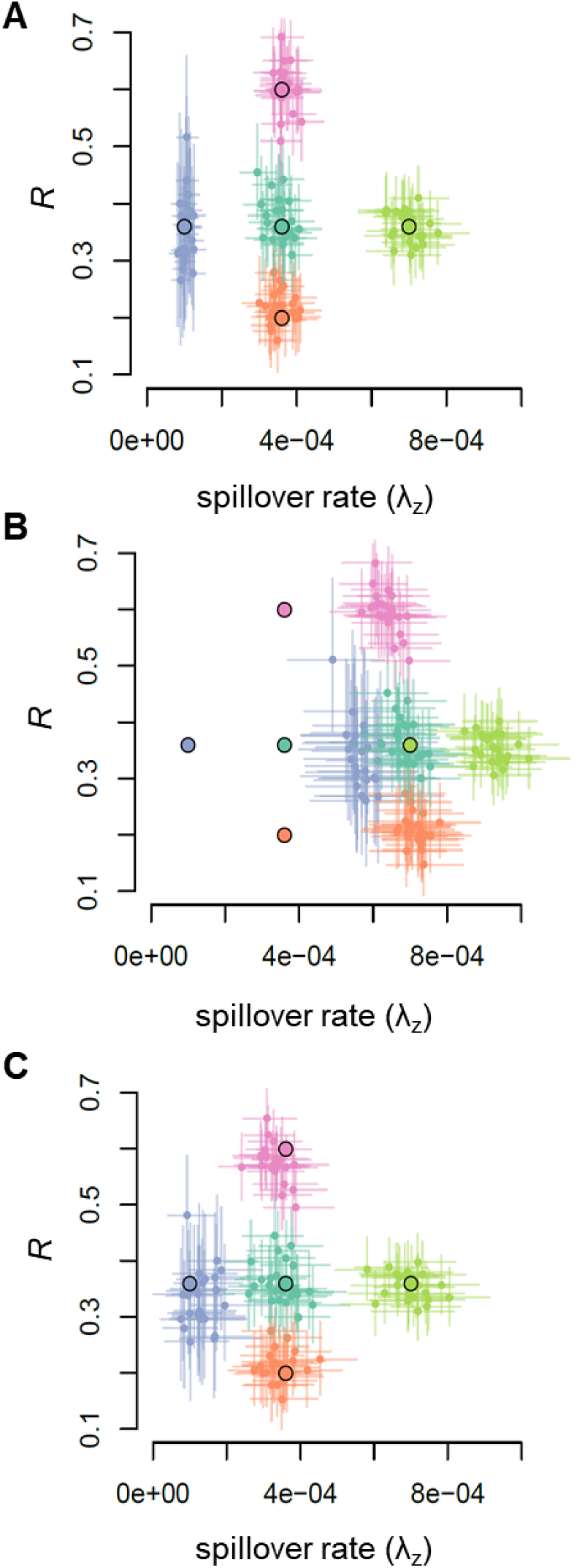
Comparison of true and inferred parameter values in simulation study. Within each color, large points outlined in black indicate the true parameter set and smaller points indicate the inferred parameter values from simulated datasets (lines show the 95% credible interval). Inferences were performed **A)** when the true number of localities under surveillance was known, **B)** when the true number was unknown and it was assumed that the number of observed localities was the total number of localities, and **C)** when the true number of localities was unknown and the corrected-denominator method was used to control for the locality observation process.

However, this method assumes that the true number of localities under surveillance is known. In real-world situations, ‘silent’ localities that experience zero cases often do not appear in the dataset, resulting in an unknown true number of surveilled localities. We investigated possible biases in parameter estimates that could arise from assuming that the number of localities that reported one or more cases represents the total number of localities under surveillance. To do so, we used the same set of simulated datasets as described above, but removed knowledge about the number of silent localities. In these datasets, silent localities make up between 21% and 85% of all localities under surveillance, with the proportion driven primarily by the spillover rate. Estimates for the reproductive number *R* were not strongly impacted (95.2% of the 95% CIs contained the true value with an average percent error of 8.4%), but the spillover rate *λ_z_* was consistently overestimated (Fig 3B). The true value for *λ_z_* was contained in none of the simulations’ 95% CIs and the posterior mean had an average percent error of 153% (S1 Table).

To further investigate the effect of this data truncation (whereby localities with zero cases do not appear in the dataset), we performed inference assuming that the observed localities represented all, 1/2, or 1/5 of the total localities under surveillance. While this assumption had a relatively small impact on the estimated *R*, it greatly impacted the inferred *λ_z_* (which is measured as the number of spillover events *per locality* per day and is therefore strongly affected by changes in the assumed number of localities) (S1 Fig). Assuming that a larger fraction of surveilled localities appear in the dataset resulted in substantially higher estimated spillover rates.

#### Corrected-denominator method (conditions on the locality observation process)

Because the total number of localities assumed to be under surveillance has a substantial impact on parameter estimates, we developed a modified version of the likelihood function that accounts for localities that were under surveillance but never observed in the dataset. This approach calculates the likelihood of the observed dataset conditional on the fact that only localities with one or more cases are included (details on the modified likelihood function can be found in Methods and S1 Text).

We tested the performance of the corrected-denominator method against simulated datasets, looking at the same parameter sets as in the first section. The inferred parameter values cluster well with their corresponding true values (Fig 3C): mean percent error in *R* estimates was 8.4% and in *λ_z_* estimates was 14.0%. Across the 125 simulations, the true parameter value was included in the 95% CI 116 times (92.8%) for *R* and 117 times (93.6%) for *λ_z_* (S1 Table).

Because an estimate of the true number of localities under surveillance would help determine the size of the population that could be detected for a given system, we assessed how well we could approximate this value. Given the number of localities with one or more cases and the mean parameter estimates, it is possible to calculate the expected total number of localities under surveillance (see S1 Text). Estimates of the true number of localities calculated for the simulated datasets center on the correct value (S2 Fig). The magnitude of estimate error is driven by the spillover rate, which largely determines the proportion of localities that are observed by surveillance. The mean percent error across simulations with spillover rate of 0.0001, 0.00036, and 0.0007 were 25.4%, 7.9%, and 2.4%, respectively, while simulations with spillover rates of 0.004 and above almost always recorded at least one case in each locality during the five year surveillance period and therefore tended to estimate the exact true number of localities.

#### Inferring the sources of transmission events

We investigated how well sampled transmission trees recovered the source of individual cases as well as higher-order measures, such as the fraction of cases originating from zoonotic, within-locality, and between-locality transmission. We tested our method using 125 simulated datasets, with 25 datasets simulated for each of five sets of true parameter values (these are the same datasets as discussed above, simulated with *R* between 0.2 and 0.6 and spillover rate between 0.0001 and 0.0007). Two hundred plausible transmission trees were sampled for each simulated dataset.

When comparing the overall fraction of cases attributed to each source type (zoonotic versus within-locality versus between-locality transmission), the sampled transmission trees closely match the true transmission patterns (Fig 4). On average, the difference between the true fraction of cases caused by zoonotic spillover and the fraction inferred in a tree was 0.022 (standard deviation 0.018), the difference for within-locality transmission was 0.006 (standard deviation 0.005), and the difference for between-locality transmission was 0.022 (standard deviation 0.018).

**Fig 4.**
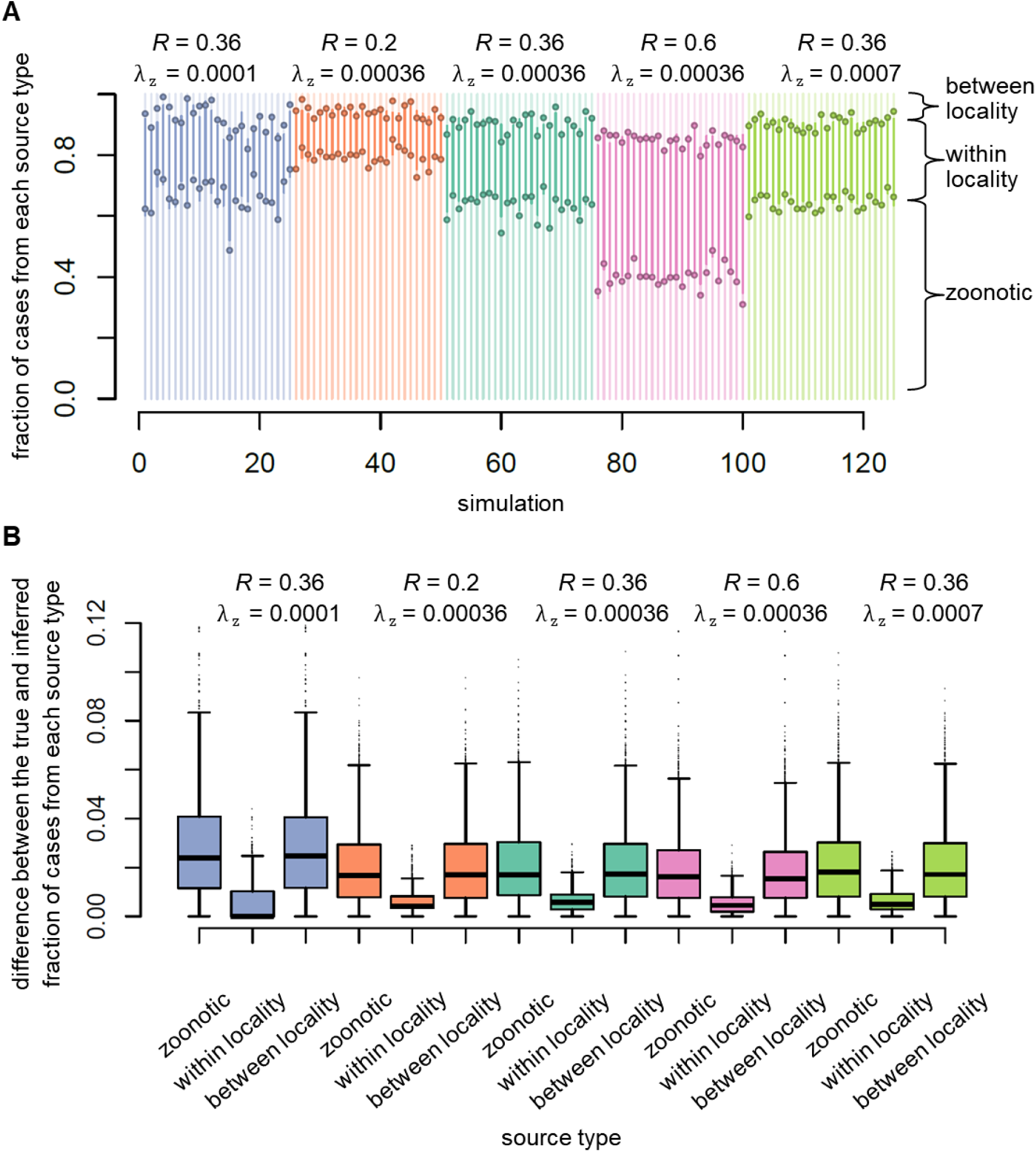
Comparison of the true and inferred fraction of transmissions from each source type. For each of five parameter sets, 25 datasets were simulated and 200 transmission trees were sampled for each of these simulated datasets. **A.** Stacked bars show the true fraction of transmissions from zoonotic (bottom bar, medium-darkness), within-locality (middle bar, light color), and between-locality (top bar, darkest color). Points on the bars indicate the inferred values. If the fraction of transmissions for each source is perfectly inferred, points will lie exactly on the transition between bar colors. **B.** Box plots summarize the error in the inferred fraction of cases originating from each source type. The error size is small across all parameter sets, especially for within-locality human-to-human transmission. The upper whisker was calculated as min(max(x), Q_3_+1.5*IQR) and the lower whisker was calculated as max(min(x),Q_1_-1.5*IQR).

The success at recovering individual transmission links was high overall but varied slightly depending on the true parameters underlying the simulation (S3 Fig). On average, sampled transmission trees inferred 85.9% of all sources correctly. Better performance was observed for lower spillover rates and lower *R*, presumably due to the fewer opportunities for misattribution of cases. Some transmission links were more likely to be captured than others: on average 90.9% and 90.1% of sampled trees correctly inferred links with zoonotic and within-locality sources, respectively, but only 36.8% of trees correctly identified the source of between-locality transmission events.

### Epidemiological insights into monkeypox

#### Applying the corrected-denominator method to 1980s monkeypox surveillance data

Between 1982 and 1986, the active monkeypox surveillance program in the Democratic Republic of the Congo detected 331 human cases in 171 localities [43]. For each human case, we know the name of the locality as well as the district or administrative subregion (henceforth referred to simply as ‘district’) and region to which it belongs. However, the total number of localities that would have been detected by surveillance had they experienced a case is unknown. We therefore used the corrected-denominator method to generate estimates under four different assumptions about which administrative unit most suitably represents the broader contact zone. The country-level, region-level, and district-level models correspond to progressively smaller choices of broader contact zones, while the locality-level model assumes that all instances of human-to-human transmission occur within a locality. We anticipate that assuming an excessively large broader contact zone could result in overestimating *R* and underestimating *λ_z_* if too many spurious human-to-human transmission events are inferred from pairs of cases that just happen to occur within a generation-time interval of one another, while assuming an inappropriately small broader contact zone could result in the opposite parameter biases if the model is unable to detect actual incidents of human-to-human transmission because the cases occur in different (assumed) broader contact zones.

In the monkeypox analysis, the size of the administrative unit used as the broader contact zone has a strong effect on the resulting parameter estimates (Fig 5A). When larger administrative units are assumed to represent the broader contact zone, a given pair of cases is more likely to belong to the same broader contact zone, giving the model more opportunities to infer inter-locality human-to-human transmission events and resulting in larger estimated reproductive number *R* and a smaller spillover rate *λ_z_*. Mean values of the posterior distribution of *R* range from 0.29 when transmission is assumed to occur only within localities to 0.52 when transmission is assumed to occur among all localities in the country (Table 1).

**Fig 5.**
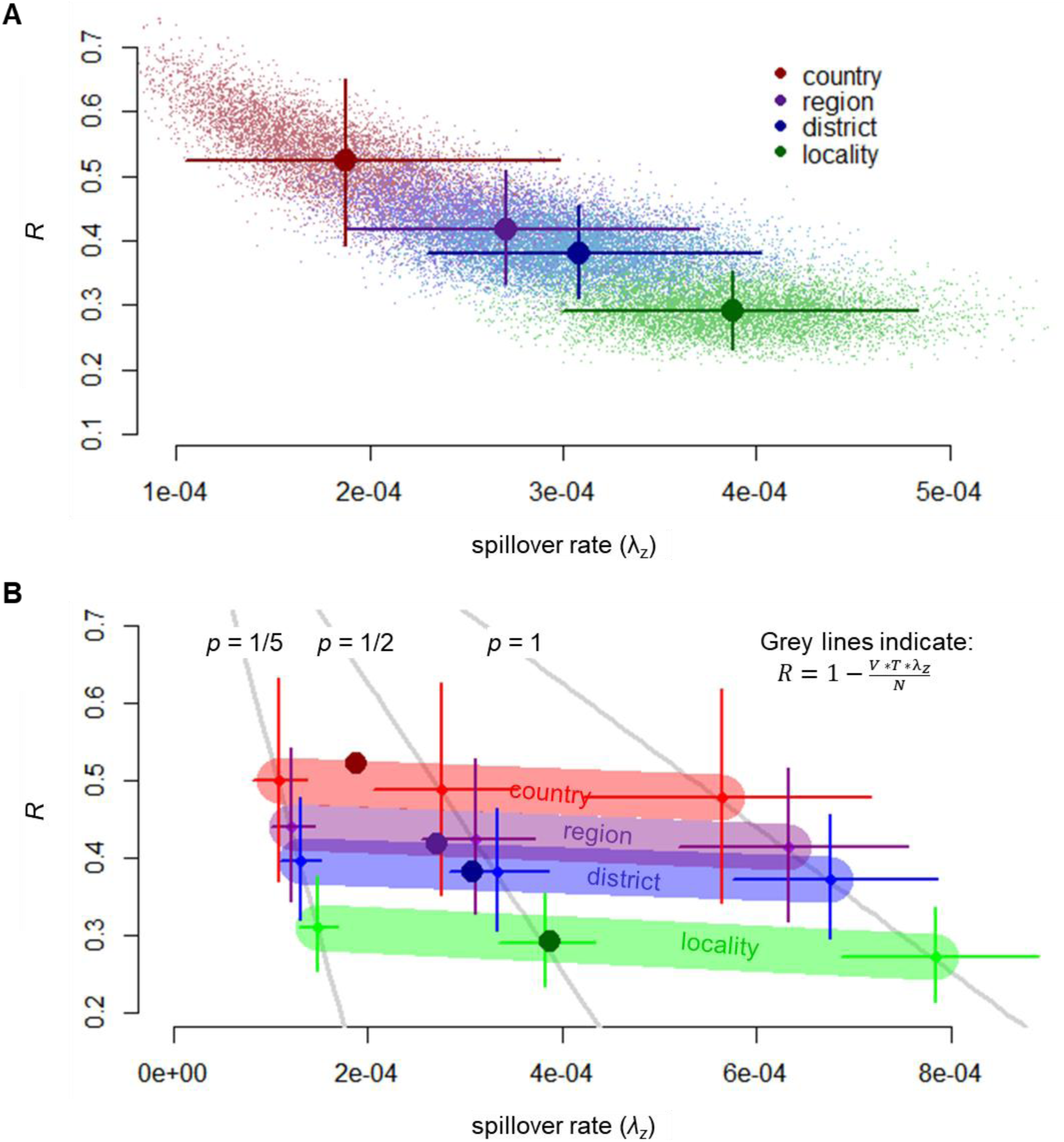
Assumptions about the broader contact zone and the total number of localities under surveillance affect parameter estimates for the monkeypox dataset. Estimates and 95% CIs for the reproductive number (*R*) and the spillover rate (λ*_z_*) of the monkeypox dataset are shown for each of the four choices of spatial scale for the broader contact zone (locality = green, district = blue, region = purple, country = red). **A.** Inference performed using the corrected-denominator method that accounts for silent localities. Light background dots are draws from the posterior, larger dots designate the mean value, and bars indicate the 95% CI. **B.** Inference performed assuming that the fraction of localities under surveillance with one or more monkeypox cases (*p*) is 1/5, 1/2, or 1. For each assumption about the total number of localities, parameter estimates fall roughly along the line 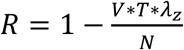 (indicated by grey lines), where *V* is the true number of localities under surveillance, *T* is the duration of surveillance, and *N* is to total number of cases. The position of estimates along this line depends on the spatial model used. Note that the slope of each line is proportional to −1/*p* because *V* = (number of observed localities) / *p*. Dots represent the mean posterior estimates and bars indicate the 95% CI. The four darker dots show the mean estimates from panel **A.**

**Table 1.**
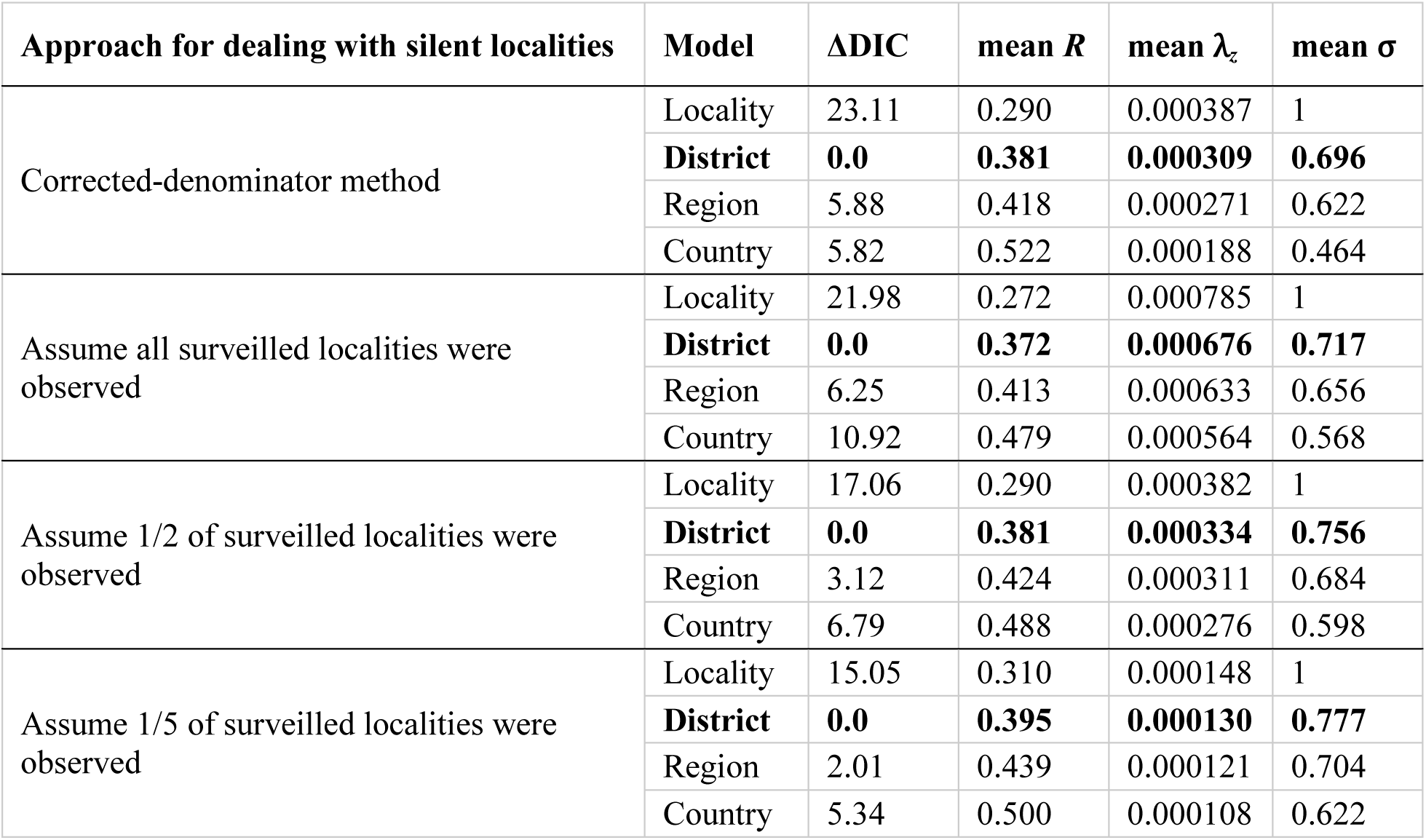
District model performs best for the monkeypox dataset in DIC model comparisons. Parameter inference for the monkeypox dataset was performed using four different approaches for dealing with the silent locality problem: the corrected-denominator method (which conditions on the observation process for localities under surveillance) and three assumptions about the fraction of localities under surveillance that were observed. For each of these approaches, inference was repeated using four choices for the broader contact zone and the DIC was calculated. Parameter estimates and ΔDIC values are shown. The model with lowest ΔDIC is preferred and is shown in bold text.

| Approach for dealing with silent localities | Model | ΔDIC | mean $R$ | mean $\lambda_z$ | mean $\sigma$ |
| --- | --- | --- | --- | --- | --- |
| Corrected-denominator method | Locality | 23.11 | 0.290 | 0.000387 | 1 |
|  | **District** | **0.0** | **0.381** | **0.000309** | **0.696** |
|  | Region | 5.88 | 0.418 | 0.000271 | 0.622 |
|  | Country | 5.82 | 0.522 | 0.000188 | 0.464 |
| Assume all surveilled localities were observed | Locality | 21.98 | 0.272 | 0.000785 | 1 |
|  | **District** | **0.0** | **0.372** | **0.000676** | **0.717** |
|  | Region | 6.25 | 0.413 | 0.000633 | 0.656 |
|  | Country | 10.92 | 0.479 | 0.000564 | 0.568 |
| Assume 1/2 of surveilled localities were observed | Locality | 17.06 | 0.290 | 0.000382 | 1 |
|  | **District** | **0.0** | **0.381** | **0.000334** | **0.756** |
|  | Region | 3.12 | 0.424 | 0.000311 | 0.684 |
|  | Country | 6.79 | 0.488 | 0.000276 | 0.598 |
| Assume 1/5 of surveilled localities were observed | Locality | 15.05 | 0.310 | 0.000148 | 1 |
|  | **District** | **0.0** | **0.395** | **0.000130** | **0.777** |
|  | Region | 2.01 | 0.439 | 0.000121 | 0.704 |
|  | Country | 5.34 | 0.500 | 0.000108 | 0.622 |

We used the mean parameter estimates obtained using each of the four broader contact zone assumptions to generate estimates of the expected total number of localities under surveillance. While only 171 localities were observed in the dataset, estimates of the total number of surveilled localities ranged from 337 (using the locality-level model) to 408 (using the country-level model). The district-level and region-level models generated similar estimates of 351 and 366 total localities, respectively.

#### Insights into how underlying assumptions drive monkeypox estimates

We investigated how different assumptions about the true number of localities and the spatial scale of human-to-human transmission would affect the parameter estimates for the monkeypox system. To explore how the presence of silent localities affects results, we repeated the analysis using the basic method (which does not account for silent localities) under the assumption that the localities observed in the monkeypox dataset represent all, 1/2, and 1/5 of the total number of localities that were under surveillance. Furthermore, for each of these assumptions about the total number of localities under surveillance, we repeated the analysis using the four different choices of broader contact zone to determine how the assumed spatial scales of inter-locality transmission impacted inference results.

Both the choice of broader contact zone and the assumed total number of localities have a large impact on estimates of *R* and *λ_z_* (Fig 5B). As noted above, models assuming smaller broader contact zones allow fewer opportunities for human-to-human transmissions to be inferred, and these models estimate substantially lower *R* values and correspondingly higher spillover rates. In contrast, assuming that a smaller fraction of surveilled localities were observed leads to slightly higher estimates of *R* and substantially lower estimates of *λ_z_* because the presence of many silent localities drives the estimate of the number of spillover events *per locality* per day lower. Estimates of *R* are most strongly affected by the choice of broader contact zone, while estimates of *λ_z_* are most strongly impacted by assumed fraction of localities observed. For all assumptions of broader contact zone and total number of localities, the means of the parameters’ posterior distributions fall along the line

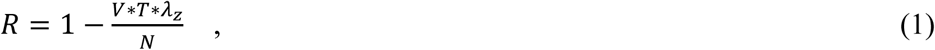

where *V* is the true number of localities under surveillance, *T* is the number of days over which surveillance occurred, and *N* is to total number of cases in the monkeypox dataset. This relationship arises because the expected number of total cases is equal to the expected number of spillover events (*V* * *T* * *λ_Z_*) multiplied by the total number of human cases expected to occur from each spillover event (1 / (1 – *R*) for 0<*R*<1). Each assumption about the total number of localities under surveillance corresponds to a separate line along which parameter estimates fall (Fig 5B). The position of the parameter estimates along this line depends on the spatio-temporal distribution of the *N* cases and the assumed spatial scale of human-to-human transmission.

#### District-level broader contact zone preferred in model comparisons

To assess which broader contact zone assumption is most appropriate for the monkeypox system, we used the deviance information criterion (DIC) to perform model comparisons for the corrected-denominator method as well as for each assumption about the number of surveilled localities. For the corrected-denominator method, the district-level model had the best DIC score, followed by the region and country-level models (Table 1). The locality-level model received a much larger DIC value, indicating that the data strongly support models that allow transmission between localities. Similarly, for each of the three assumptions about the true number of surveilled localities, the district-scale model performed best in DIC model comparisons (Table 1).

#### Inferring the sources and distances of transmission events

We used the district-level corrected-denominator method to sample 20,000 transmission trees for the monkeypox dataset. The sampled transmission trees attributed an average of 60.8% (standard deviation of 2.2%) of cases to zoonotic spillover, 28.5% (standard deviation of 0.9%) of cases to within-locality human-to-human transmission, and 10.7% (standard deviation of 2.1%) of cases to between-locality human-to-human transmission. For comparison, the results using the three other broader contact zone assumptions are shown in S4A Fig. Each model’s trees include a similar number of within-locality human-to-human transmission events, but increasing the spatial scale of the broader contact zone increases the number of inferred between-locality transmission events.

To characterize the distance range over which inter-locality transmission occurs, we focused on links in the sampled transmission trees that occurred between cases with known GPS coordinates (280 out of 331 monkeypox cases had recorded GPS coordinates). The number of transmission events in each sampled tree that occurred over a certain distance was then compared to the number of transmission events expected to occur over each distance if transmission between all localities in a broader contact zone was equally likely (see Methods 4.3 for how this ‘null distribution’ was calculated).

For all models allowing inter-locality transmission, more transmission events were inferred to occur across ≤ 30 kilometers than expected based on the null distribution (Fig 6, S4B Fig). For each inferred transmission tree, a binomial test was used to examine whether more transmissions were inferred to occur over ≤ 30 kilometers than expected based on the null distribution of transmission distances. Out of 20,000 sampled trees for each model, p-values of less than 0.1 were obtained in 93% of the district, 72% of the region, and 81% of the country-level models’ trees. The median p-values for these three models were 0.007, 0.030, and 0.012, respectively (S5 Fig shows the full distributions of p-values obtained across all sampled trees).

**Fig 6.**
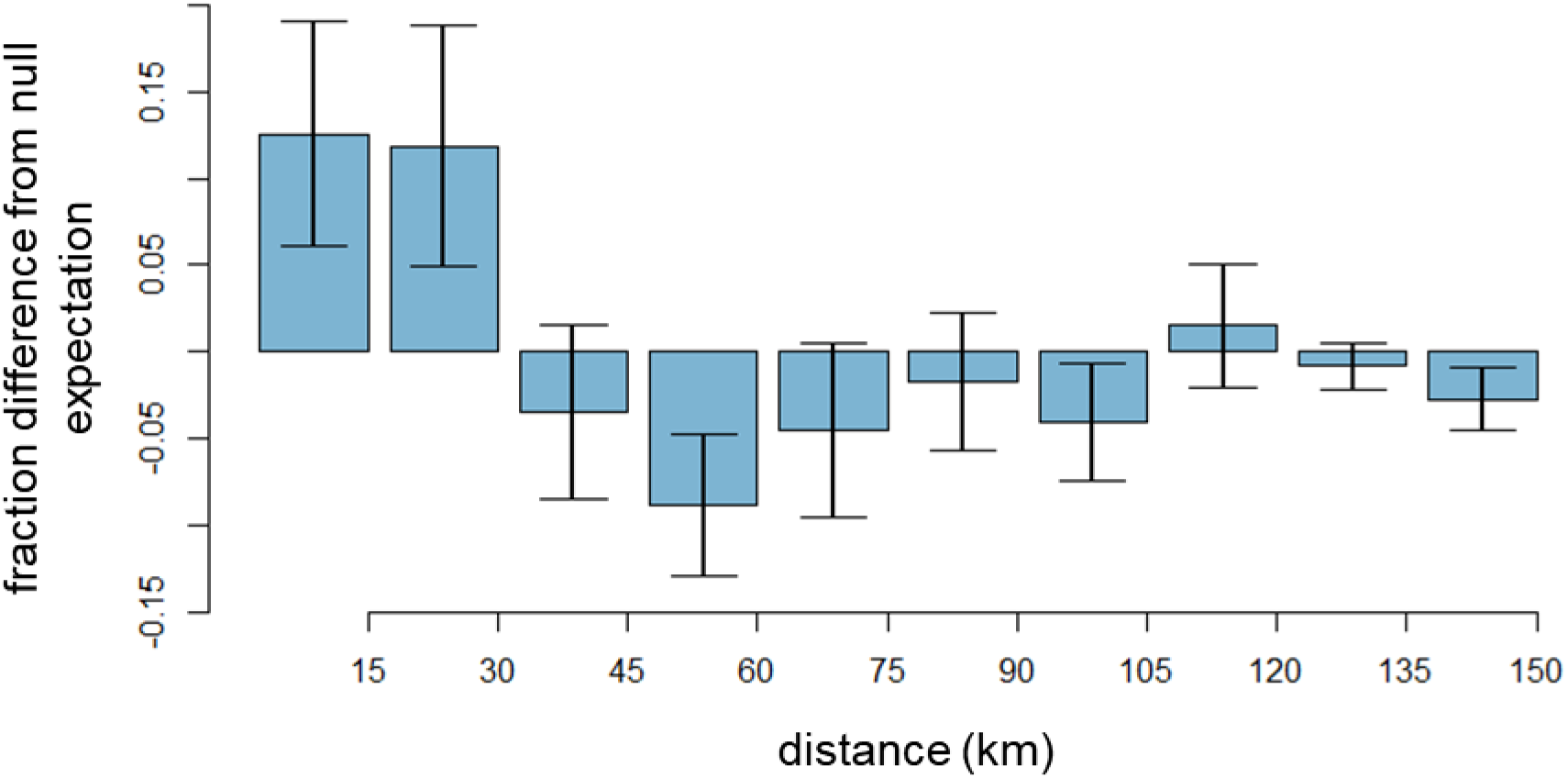
Distance of inferred inter-locality human-to-human transmission events. Shaded bars show the difference between the mean proportion of inter-locality human-to-human transmissions inferred to occur over a given distance by the district model and the proportion expected based on the spatial distribution of localities (the ‘null expectation’). Error bars show the standard deviation among all inferred transmission trees.

#### Comparison of sampled transmission trees with contact-tracing data

Contact-tracing, where the contacts of a case were recorded and follow-up investigations determined whether or not the contacts had become infected, was done for a subset of monkeypox cases. Instances where a contact developed an infection are presumed to be instances of human-to-human transmission. For each of these epidemiologically contact-traced links, we looked at how frequently the sampled transmission trees for each model captured the transmission link.

Of the 53 case pairs linked through contact tracing, an average of 79.5% (standard deviation of 4.2%) were recovered in each of the district model’s sampled transmission trees (Fig 7A). The highest success was seen for pairs of epidemiologically-linked cases whose dates of symptom onset were between 7 and 25 days apart (Fig 7B). Although it is generally believed that the generation interval for human-to-human transmission of monkeypox is between 7 and 23 days [43,53], several case pairs that occurred more than 23 days apart were epidemiologically linked through contact-tracing. It is possible that these links, which were often missed in the sampled transmission trees, are not true instances of human-to-human transmission. Cases that occurred in different localities were also less likely to be linked in a sampled transmission tree, though even for these inter-locality pairs, the district-level model tended to perform better than the other three models (S6 Fig). The four models had similar success at recovering within-locality links. In all models, when a link was incorrectly inferred, it frequently was inferred to originate from zoonotic spillover instead. Although the district model had the highest success at recovering contact-traced links, the sampled trees from all models recovered an average of >76% of contact pairs.

**Fig 7.**
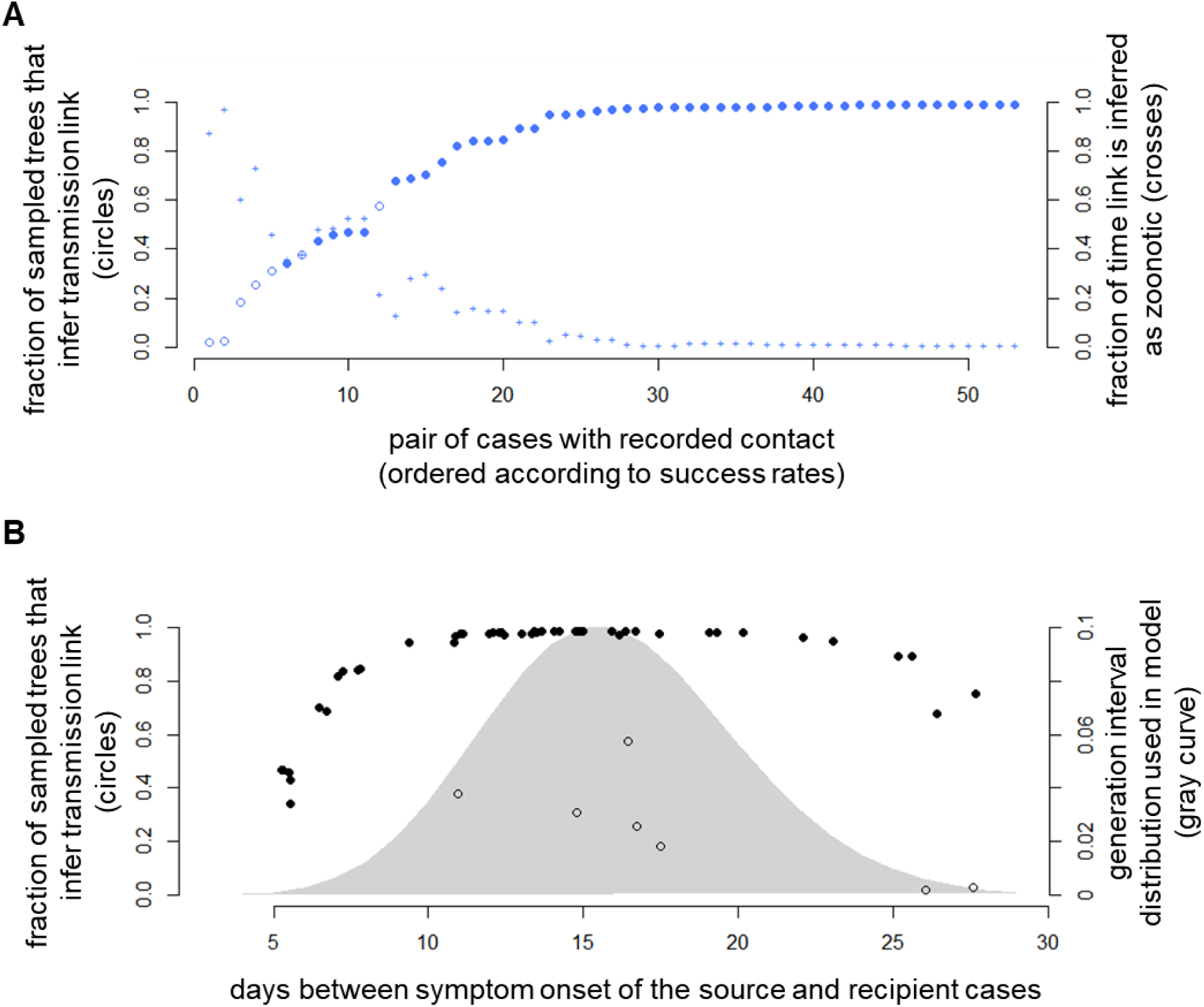
Comparison of epidemiologically contact-traced links with sampled transmission trees. **A.** <u>Circles</u> (left axis) show the fraction of sampled trees that infer the epidemiologically-traced source. Open circles represent inter-locality links while closed circles represent intra-locality links. <u>Crosses</u> (right axis) indicate the probability that a link is instead inferred to have a zoonotic source. Results are shown for the model assuming the district-level broader contact zone. Links are sorted from lowest to highest success. **B.** The fraction of sampled transmission trees that recover a contact-traced link is influenced by the number of days between the symptom onset of source and recipient cases. <u>Circles</u> (left axis) show how often a given link was inferred as a function of the generation interval while the <u>gray curve</u> (right axis) shows the probability density for the generation interval assumed by the model.

Comparison of the transmission tree generated using only contact-tracing data with the trees created using the district-level and locality-level models highlights how much our perception of the transmission dynamics depends on assumptions about spatial spread (Fig 8). Most of the within-locality transmission links detected through epidemiological contact-tracing appear in the locality-level model’s tree, though the locality-level tree suggests substantially more human-to-human transmission events than captured in the contact-tracing tree. However, the locality-level tree misses all inter-locality links. The district-level model’s tree captures most of the links indicated by the locality-level tree, and also suggests that inter-locality transmission is occurring, though it has low power to determine exactly which case pairs are linked through inter-locality transmission.

**Fig 8.**
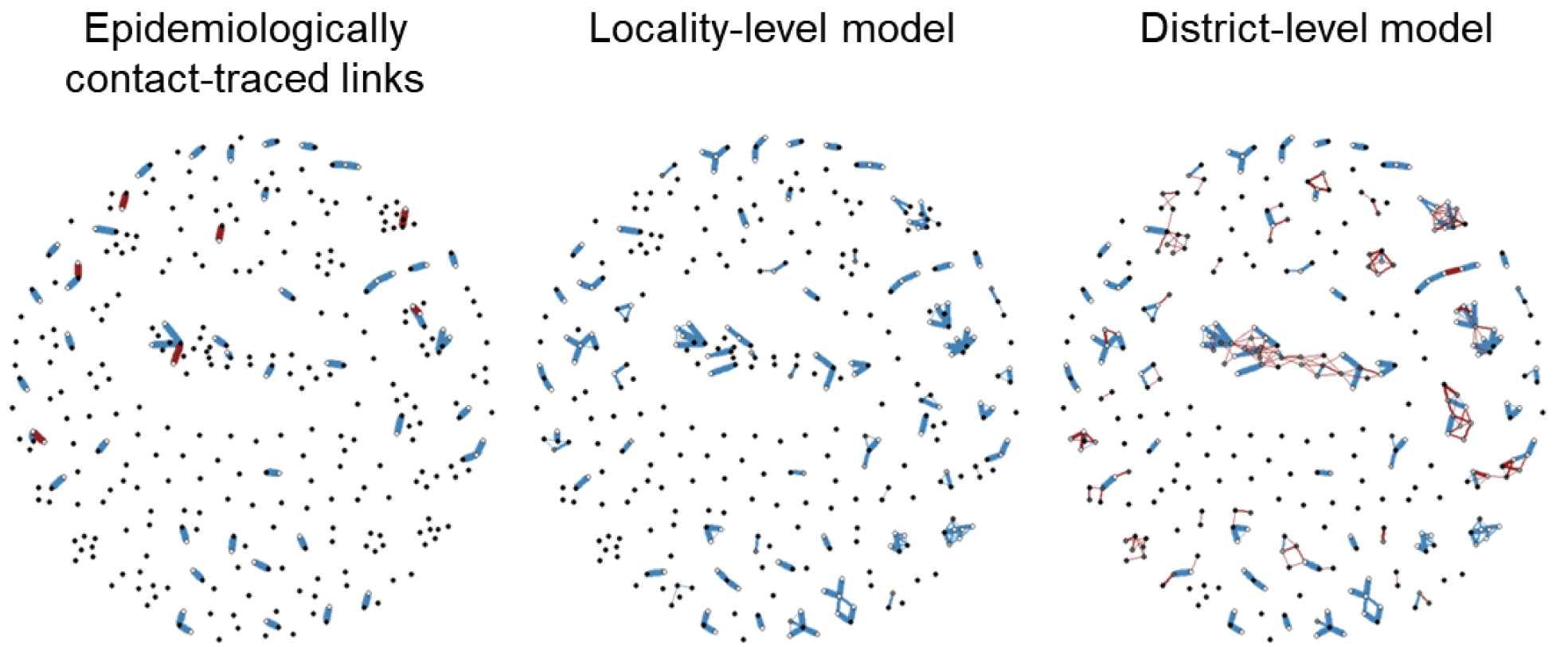
Comparison of monkeypox transmission trees created from contact-tracing, the locality-level model, and the district-level model. Points represent cases and edges indicate inferred transmission links between cases. Edge thickness corresponds to the frequency with which a given transmission link was inferred while edge color indicates whether a pair of linked cases occurred within the same (blue) or different (red) localities. The darkness of a point’s fill indicates how frequently the case was inferred to have a zoonotic source, so transmission links often go from black points (cases caused by zoonotic spillover) to white points (cases infected by a human source).

### Sensitivity analyses

We conducted a variety of sensitivity analysis tests using simulated datasets to assess how robust the method was over a range of parameter values and assumption violations (full descriptions are provided in S1 Text). The method continued to perform well even at very high spillover rates (S7 Fig, S2 Table) and when the offspring distribution used in simulations differed from the one assumed in the inference (S8 Fig, S3 Table). In some situations, assuming a larger broader contact zone than the one used for simulations could lead to an overestimation of *R* and an underestimation of *λ_z_* (S4 and S5 Tables). This outcome is consistent with what was observed in the monkeypox analysis where assuming a larger spatial scale for the broader contact zone corresponded to a higher estimate of *R* and a smaller estimate of the spillover rate (Fig 5). When simulations were run with highly structured, non-homogeneous spillover, substantial biases in the inference results occurred because this spillover process gives rise to clusters of primary cases that the model mistakes as arising from human-to-human transmission (S9 Fig).

## Discussion

### Principal findings

In this work, we developed and tested a method to infer fundamental epidemiological parameters and transmission patterns for zoonotic pathogens from epidemiological surveillance data with aggregated spatial information. When tested against simulated datasets, the method successfully recovered estimates of *R* and spillover rate close to the true values and also inferred the fraction of cases resulting from zoonotic, within-locality, and between-locality sources with a high degree of accuracy. The ‘unknown denominator problem’ that occurs when the total number of localities under surveillance is unknown can cause large biases in parameter estimates, so we modified the inference method to account for this observational process and enable unbiased estimation in the presence of this common data gap.

We applied the method to a rich surveillance dataset of human monkeypox in the Congo basin from the 1980s and found that human-to-human transmission of monkeypox between localities plays an important role in the pathogen’s spread. Of the four assumptions we tested for the spatial scale of the broader contact zone, the district-level model was best supported by DIC model comparisons and validation with contact-tracing. In addition, the signal of elevated inter-locality transmission occurring over ≤ 30 kilometers suggests that most inter-locality transmissions occur in a relatively small neighborhood, consistent with the limited transportation infrastructure in the DRC. This further corroborates that the district-level model, which is the smallest spatial aggregation scale that still permits inter-locality transmission, is likely the most appropriate choice for capturing inter-locality transmission patterns of human monkeypox.

The district-level model estimates a reproductive number for human monkeypox of 0.38 (0.31-0.45 95% CI). This value is slightly higher than previous estimates of *R* for the 1980s DRC monkeypox dataset, which was estimated as 0.30 (90% CI 0.22-0.40) in Blumberg and Lloyd-Smith [16], as 0.32 (90% CI 0.22-0.40) in Lloyd-Smith et al. [54], and as 0.28 in Jezek et al. [52]. There are several explanations for the higher estimate we obtained. The previous studies may have underestimated the reproductive number, particularly if contact-tracing or cluster formation methods were liable to miss transmissions that occurred between localities. Indeed, the estimate obtained using the locality-level model (*R* = 0.29) closely matches previous estimates. It is also possible that the district-level model may overestimate the amount of human-to-human transmission in the same way that the region- and country-level models picked up a higher signal of human-to-human transmission than the district-level model due to their larger broader contact zone sizes. The size of the DRC’s districts and administrative subregions used for the district-level model vary in size, but average around fifteen thousand square kilometers, or around one hundred forty kilometers across, encompassing a much greater distance than most human-to-human transmission events likely occur over. We therefore expect that the true value of *R* is bounded by the estimates of the locality-level and the district-level models.

In addition to providing an estimate of monkeypox’s reproductive number, the methods give insight into the frequency of spillover and the spatial scale of human-to-human transmission. The district-level model estimates a mean spillover rate of around 0.11 spillover events per locality per year, which corresponds to roughly one spillover event every nine years in each locality. It also estimated that around 70% of human-to-human transmissions occur within a locality. This finding contrasts with the assumption that human-to-human transmission occurs within a locality, which is commonly used to generate transmission clusters, and suggests that estimates generated using that assumption may substantially underestimate the amount of human-to-human transmission occurring in the system. The importance of inter-locality contacts has been reported for the neighboring country of Uganda, where a survey by le Polain de Waroux et al. [55] on rural movement and social contact patterns indicated that 12% of social contacts occurred outside participants’ village of residence.

Among human monkeypox cases with recorded geographical coordinates, a clear signal emerged of higher rates of human-to-human transmission between localities ≤ 30 kilometers apart. This pattern seems reasonable given the infrastructure and general difficulty of transportation in the more remote regions of the DRC. It also suggests a similar pattern of movement as found in the le Polain de Waroux et al. [55] survey. Their analyses indicate that 90% of people who traveled outside their village of residence remained within 12 km.

### Spatial scale of transmission and aggregated spatial data

The potential biases introduced when analyzing data reported at a course spatial scale have been explored in a wide range of contexts [56–58], yet the implications of using this type of spatial information to infer the transmission dynamics of an infectious disease is not obvious. When spatial information is only reported at the level of large spatial zones like districts, regions, or countries, no finer-scale information is available to inform which human cases transmitted infection to one another between different localities. Here we explored how the size of these spatial zones would affect inference for the monkeypox system by repeating the analysis using spatial information at the district, region, or country resolution. The large differences in parameter estimates generated under different broader contact zone assumptions in the monkeypox analysis illustrates how sensitive inference results can be to the spatial scale assumed for human-to-human transmission, and suggests that reporting spatial data at too large a scale or ignoring inter-locality transmissions can lead to substantial estimate biases.

In the context of monkeypox in the DRC, analysis of simulations using the exact geographic coordinates reported for 80% of localities in the monkeypox surveillance dataset replicated the increasing estimates of *R* and decreasing estimates of spillover rate as the spatial aggregation scale increased (S4 and S5 Tables). However, the magnitude of the effect in simulated datasets was smaller than in the monkeypox analysis. This could be a result of the particular assumptions about inter-locality transmission patterns used in the simulations, but it also opens the question of whether outside large-scale factors such as seasonality or fluctuations in surveillance effort might induce temporal autocorrelation among unlinked human cases, giving rise to temporal clustering of cases that the model interprets as human-to-human transmission.

This analysis serves to emphasize the importance of selecting an appropriate spatial scale and using caution when interpreting results obtained using spatially aggregated data. Many methods implicitly assume a certain scale of spatial transmission, often ignoring the possibility of longer-range transmissions, so careful consideration of whether that scale is appropriate for the system is essential.

In general, recording precise spatial locations of cases is vital for increasing the inferential power of modeling analyses. Developing methods that maintain spatial information without risking a breach in confidentiality is a nontrivial challenge, but progress has already been made in generating possible solutions such as geographic masking or the verified neighbor approach [59,60].

### Model assumptions and future directions

In this work, we assumed that the spillover rate was homogenous through time and space, but more complex disease dynamics in the reservoir or spatiotemporal heterogeneity in animal-human contacts may cause nontrivial deviations from this assumption in real-world systems. Of particular concern is the possibility that outbreaks in the reservoir could cause periods of amplified local spillover, which could create a clustering pattern of human cases potentially indistinguishable from human-to-human transmission. Without information about disease dynamics in the reservoir, accounting for this heterogeneous spillover will be challenging, but certain types of pathogen dynamics, such as seasonal epidemics or expanding wave-fronts of infection, could be incorporated into the model structure.

Similarly, spatially and temporally variable surveillance intensity could potentially mimic the signal of human-to-human transmission clusters and result in overestimates of the reproductive number. Future surveillance programs could help mitigate this challenge by recording a measure of surveillance effort undertaken at each location and time.

This work assumes that *R* is constant across all localities; however, to obtain a full picture of pathogen emergence risk, it may be necessary to consider the heterogeneity in transmission intensity among different human populations, as well as the interplay between where *R* is highest versus where spillover tends to occur [61]. In some zoonotic systems, for instance, spillover predominantly occurs into remote villages and towns that are in close proximity to forested regions. However, we generally expect these villages to have lower levels of human-to-human transmission than the more densely-packed cities [62–64]. A pathogen may even be incapable of supercritical spread until it reaches such a city. Therefore, to assess the probability a pathogen will successfully emerge and to determine which populations to target with control measures, it may be necessary to establish not only the spillover rate and *R* across different populations, but also the rate of dispersal of the pathogen between those populations [61].

Several assumptions may need to be modified when applying this method to other zoonotic systems. Because we assume that the source of human-to-human transmission events will show symptoms before the recipient, the likelihood function can treat human cases as occurring independently conditional on preceding cases. For zoonotic diseases in which infected individuals frequently transmit the pathogen before showing symptoms (or when asymptomatic cases contribute non-negligibly to transmission), the likelihood expression would need to be modified substantially, and the lack of independence between cases might make a simulation-based inference approach necessary.

We assume that sufficiently few infections occur relative to the population size that depletion of susceptible individuals does not affect transmission dynamics. While appropriate when there are few human infections or in the early stages of invasion, this assumption could bias estimates if applied in a system with sufficiently high levels of human infection or where transmission occurs primarily among highly clustered contacts, such as individuals within a household. We also note that in the monkeypox example we are estimating the *effective* reproductive number, which takes into account existing population immunity. If the goal instead were to establish the basic reproductive number (the reproductive number for the pathogen in a fully susceptible human population), accounting for past exposure to the pathogen or other cross-immunizing pathogens or vaccines would be necessary.

The current methods assume that all human cases that occur during the surveillance period inside the surveillance area are observed. This assumption is plausible for the analysis of the 1980s monkeypox dataset, given the unusually high resources and experience level of this surveillance effort in the aftermath of the smallpox eradication program and the use of serology to detect missed cases retrospectively [43]. However, most zoonotic surveillance systems operate with limited resources and have a much lower detection rate. Ignoring unobserved cases will lead to underestimation of the spillover rate, while the effect on estimation of *R* will depend on the nature of the surveillance program. For instance, in the chain-size analyses of Ferguson et al. [28] and Blumberg and Lloyd-Smith [16], *R* is underestimated when the detection probability of each case is independent of one another or when right-censoring occurs but overestimated when a detected case triggers a retrospective investigation that detects all cases in that transmission chain.

### Conclusions

This work expands our ability to assess and quantify important zoonotic pathogen traits from commonly available epidemiological surveillance data, even in the absence of exact spatial information or a complete count of localities under surveillance. We anticipate that these methods will have greatest value in the common circumstance when the source of cases, particularly whether a case came from an animal or human source, cannot be readily established. In such situations, the ability to infer the pathogen’s reproductive number, spillover rate, and spatial spread patterns from available surveillance data, will greatly enhance our understanding of the pathogen’s behavior and could provide valuable insights to help guide surveillance design and outbreak response.

## Methods

### Model

In broad terms, the model describes the probability of observing a set of symptom onset times and locations of human cases given the timing and location of previous cases and parameters that underlie the transmission process. Human infections can arise from either animal-to-human transmission (‘zoonotic spillover’) or human-to-human transmission (Fig 1B). Human-to-human contact occurs more frequently within a locality than between localities, but can still occur between localities that belong to the same broader contact zone (Fig 1A).

All sources of infection are assumed to generate new cases independently of one another. The number of human cases that become symptomatic on each day in each locality caused by zoonotic spillover is assumed to follow a Poisson distribution with mean *λ_z_*. For simplicity and because reservoir disease dynamics are rarely well characterized, we assume the Poisson process is homogenous through time and across localities, but this assumption could be modified for a system where more information is available about the reservoir dynamics (e.g., [34]). New infections can also arise from contact with infected humans. The number of new infections that become symptomatic on day *t* in locality *v* caused by an infectious individual who became symptomatic on day *s* in locality *w* is assumed to be a Poisson-distributed random variable with mean *λ_{s,w},{t,v}_*.

Aggregating cases caused by all sources of infection (both human and zoonotic), the total number of new cases on day *t* in locality *v* is a Poisson-distributed random variable with mean

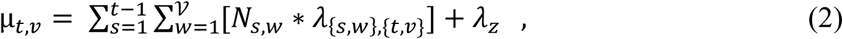

where 𝒱 is the number of localities under surveillance and *N_s,w_* is the number of cases with symptom onset on day *s* in locality *w*.

The mean of the Poisson random variable describing human-to-human transmission, *λ_{s,w},{t,v}_*, depends on the reproductive number of the pathogen in humans, the generation time distribution, and the coupling between localities:

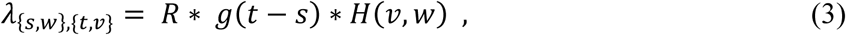

where *R* is the reproductive number of the pathogen; *g*(*t-s*) is the generation time distribution, which gives the probability that a secondary case becomes symptomatic *t-s* days after the index case shows symptoms; and *H*(*v,w*) describes the amount of transmission between localities *v* and *w* and takes values between zero (if no transmission can occur between localities *v* and *w*) and one (if all cases arising from an infected individual in locality *v* arise in locality *w*). The generation time *g*(*t-s*) is assumed to follow a negative binomial distribution. For this study, we used a mean of 16 days and a dispersion parameter of 728.7 (calculated by fitting a negative binomial distribution to observed generation interval counts for smallpox presented in Fig. 2b of [65]), which is consistent with previous estimates of the generation time for both smallpox and monkeypox [43,53,65,66].

The factor that describes the amount of transmission that occurs between localities *v* and *w* (*H*(*v,w*)) could reflect Euclidean distance, travel time, inclusion in different spatial zones, or any other available measurement. To accommodate the imperfect spatial information available for many zoonotic surveillance systems, this study focused on developing methods for the situation when only a locality name and an aggregated spatial zone (such as district or country) is reported for cases, rather than an exact position. We assume that inter-locality transmission occurs only among localities within the same broader contact zone (Fig 1A). Because transmission will be greater within a locality than between localities, a proportion *σ* of secondary cases are assumed to occur in the same locality as the source case and a proportion (*1-σ*) of secondary cases are assumed to occur amongst the outside localities that are within the same broader contact zone as the source case. This outside transmission is assumed to be divided equally among all localities within the index case’s broader contact zone:

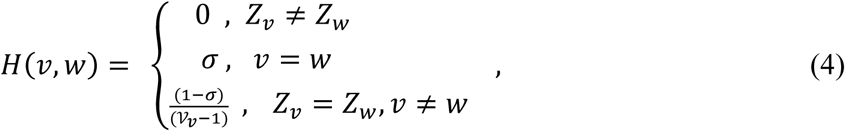

where *Z_v_* indicates the broader contact zone of locality *v* and 𝒱*_v_* is the total number of localities in the broader contact zone of locality *v*. For a given locality *v*, the sum of *H*(*v,w*) across all *w* equals one. To observe the effect of assuming different broader contact zones, the monkeypox case study was repeated under four different assumptions about the spatial scale of human-to-human transmission: locality, district, region, and country-level.

### Model inference

#### Likelihood function

Using the model described above, a likelihood function was used to evaluate a parameter set (*θ =* {*R, λ_z_, σ*}) given the data (*D* = *N_t,v_* cases observed on each day *t* and locality *v*):

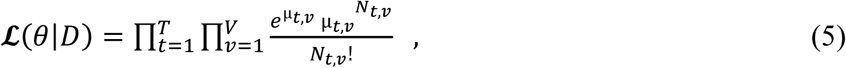

where *T* is the number of days surveillance was conducted and *V* is the total number of localities under surveillance.

While this approach works well when the total number of surveilled localities is known (see Fig 3A), localities often only appear in the dataset if they have reported cases; as a result we may not know the total number of localities under surveillance. Ignoring localities with zero cases can lead to biased parameter estimates (see Fig 3B). We explored several alternative approaches to account for these silent localities; the preferred approach rescales the likelihood function to reflect that localities with zero cases are not included in the data. Several approximations are made in this approach to estimate unknown parameters and improve computational tractability. The details of the derivation for the model are given in S1 Text, and the final likelihood function is:

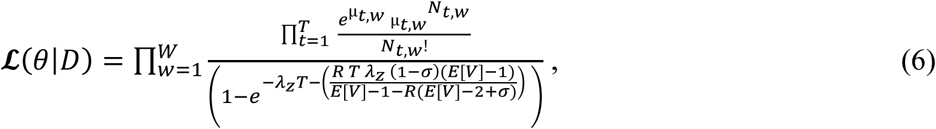

where *W* is the number of observed localities (localities with one or more cases) and *E*[*V*] is the expected number of localities given the parameter values and the number of observed localities.

#### Parameter estimation

Markov chain Monte Carlo (MCMC) was used to obtain the posterior distributions of the model parameters. The fraction of transmissions occurring within a locality (*σ*) and the reproductive number (*R*) were given uniform priors on zero to one. The expected number of spillover events per locality per day (*λ_z_*) was given a uniform prior with a lower bound of zero and an upper bound selected to be far above the converged posterior distribution (ranging from 0.0075 to 1, see S10 Fig for comparison of spillover priors and posterior distributions).

The chains were run for 100,000 steps, with a burn-in of 20,000. They satisfied visual inspection for convergence. In addition, the Gelman and Rubin multiple sequence diagnostic was evaluated for three parallel chains from each of the models for the monkeypox dataset [67]. The Gelman-Rubin potential scale reduction values were less than 1.00033 across all models, indicating that the chains have converged close to the target distribution [68].

### DIC model comparisons

For the monkeypox dataset, four assumptions about the choice of broader contact zone were compared using the deviance information criterion (DIC). This approach combines a complexity measure, used to capture the effective number of parameters in each model, with a measure of fit in order to perform model comparisons. Models are rewarded for better ‘goodness-of-fit’ to the data and penalized for increasing model complexity. Similarly to the well-known Akaike information criterion (AIC) model comparisons, models with smaller DIC values are preferred. As a rule of thumb, a difference between models’ scores of four or more generally indicates that the model with the larger value is ‘considerably less’ well supported by the empirical evidence [69]. The values necessary to calculate the DIC can be readily obtained from the MCMC output [70].

### Transmission tree reconstruction

The origin of cases (zoonotic spillover, intra-locality human-to-human transmission, or inter-locality human-to-human transmission) and the distances of inter-locality human-to-human transmission events (when case localities are known) can be established given a particular transmission tree. To gain estimates of these measures, trees were sampled based on the model and the parameter posterior distributions. From the MCMC output (representing draws from the posterior distribution), *d*_1_ sets of parameter estimates were drawn to create *d*_1_ transmission-probability matrices (***P***). The entry *P_ij_* describes the probability that individual *i* was infected by individual *j*. The diagonal values of the matrix represent the probability a case originated from zoonotic spillover. For a case *i* observed to occur on day *t* in locality *v*, the probability that case *j* was the source of case *i* (*P_ij_*) was taken to be the proportion of *µ_t,v_* (the expected total number of cases on that day and locality; defined in equation 2) contributed by case *j*. By sampling *d*_2_ transmission trees from each of these transmission-probability matrices, we calculated the proportion of cases that resulted from spillover, within-locality transmission, and between-locality transmission in each sampled tree. When testing the method using 125 simulated datasets, 200 sampled transmission trees were generated for each dataset, with *d*_1_ =20 and *d*_2_ =10. For the monkeypox dataset, 20,000 transmission trees were generated with *d*_1_ =200 and *d*_2_ =100.

For inferred inter-locality human-to-human transmission events in the monkeypox dataset, if the GPS coordinates were known for both localities in a transmission pair, the transmission distance was calculated using the *gdist* function in the R package Imap [71]. The ‘null distribution,’ used for comparing the number of inferred inter-locality transmission events with the number expected to occur if spatial location played no role in transmission, was calculated by pooling all cases for which locality GPS coordinates are known, sampling all inter-locality pairs permitted by the model, and recording the distance between the localities in each pair.

### Simulation of test datasets

To test the effectiveness of the methods, datasets with known parameter values were simulated using the model explained above. Simulations were run over 1825 days (approximately 5 years) and 325 surveilled localities. The localities were assumed to be partitioned across thirty districts and six regions, with the distribution of localities across districts and regions similar to that observed for the monkeypox dataset. The generation time interval (the number of days between symptom onset of the source and recipient cases) was assumed to follow a negative binomial distribution with a mean of 16 days and a dispersion parameter of 728.7 (as described above), with a maximum generation time interval of 40 days. A number of parameter sets, as well as different underlying model structures, were used for simulations (S6 Table). Simulation parameters were chosen to approximate the monkeypox dataset, with *σ* set at 0.75, *R* ranging from 0.2 to 0.6, and *λ_z_* ranging from 0.0001 to 0.1. Unless otherwise specified, simulations were performed assuming the district-level model. Details on the models used for sensitivity analyses that use the exact spatial location of cases or allow highly structured and non-homogenous spillover patterns are provided in S1 Text.

### Monkeypox data

Data on human monkeypox cases in the Democratic Republic of the Congo (DRC), formerly ‘Zaire,’ were collected as part of an intensive surveillance program supported by the World Health Organization. During the peak surveillance period, between 1982 and 1986 [72], data on 331 cases of laboratory-confirmed human monkeypox were recorded (see Fig 2, S1 Data) [43]. As part of field investigations, mobile teams visited the locality of a monkeypox case to collect information about the case, such as the date of fever and rash onset (for this study, the symptom onset date was taken to be the fever onset date; if the date of onset was not recorded, the rash onset date was used instead), as well as to identify individuals who had had close contact with the case [52,73]. If one of these contacts developed monkeypox within 7 to 21 days of first exposure, the presumptive source case was recorded (S2 Data) [43,73].

Between 1982 and 1986, human monkeypox cases were observed in 171 distinct localities, distributed among 30 districts and administrative subregions (simply referred to as ‘districts’) and 6 regions. The total number of localities that could have been detected by surveillance is unknown. Of the 171 observed localities, GPS coordinates are available for 136 localities (which corresponds to 280 out of 331 cases). The district, region, and country of a locality were always recorded.

## Supporting information captions

**S1 Text. Additional information on methods.** Supplementary text describing the corrected-denominator likelihood, the estimation of the total number of localities under surveillance, the simulation methods, and the sensitivity analyses.

**S1 Fig.**
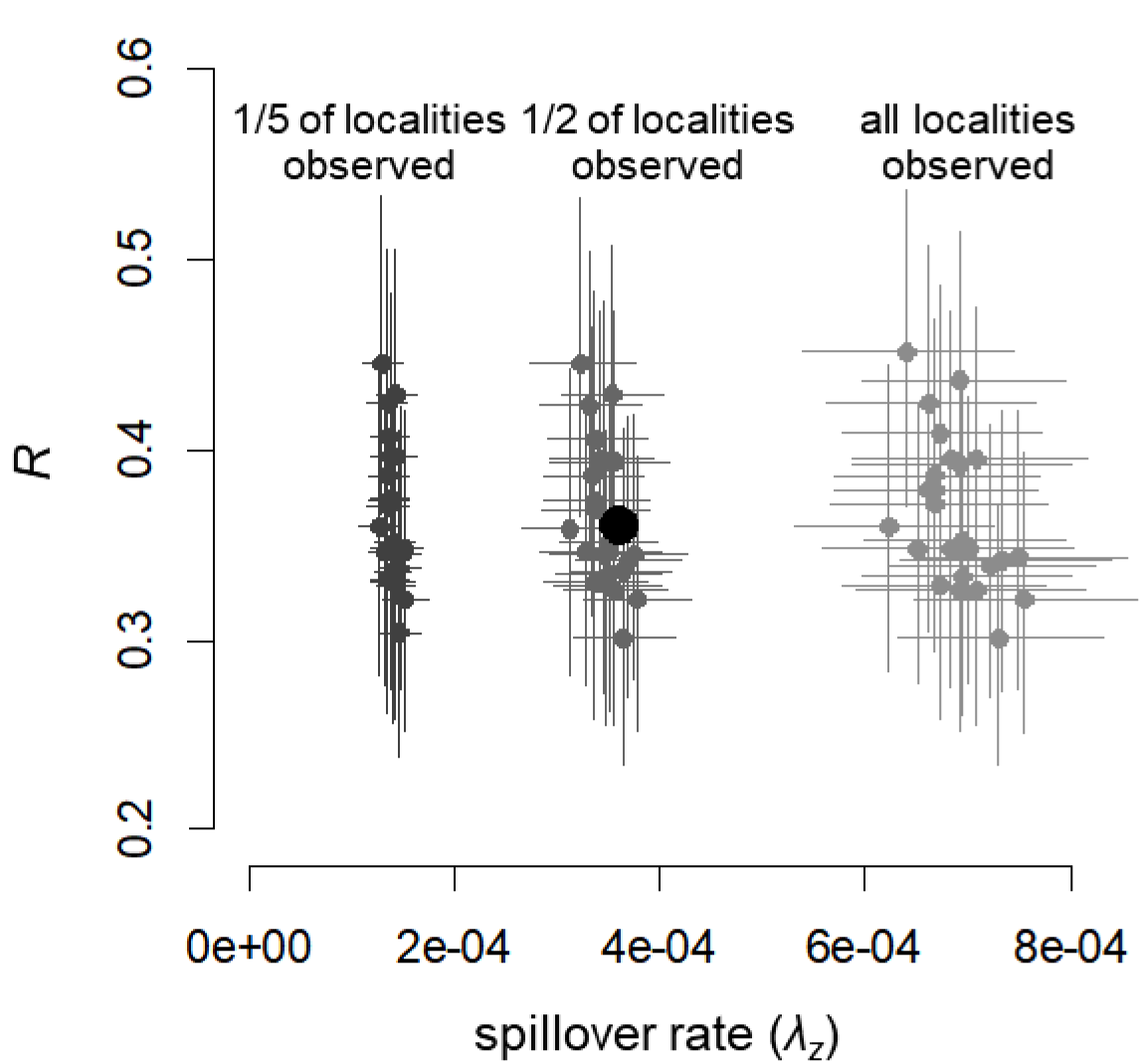
Effect of assumed fraction of localities observed on parameter estimates. The true parameter values are indicated by a large black dot and while smaller points indicate the inferred values from 25 simulated datasets (lines show the 95% credible interval). For each dataset, inference was performed assuming that 1/5, 1/2, and all of the localities under surveillance were observed. For these simulations, the true percentage of localities observed ranged from 46% to 57%, with a mean of 52%.

**S2 Fig.**
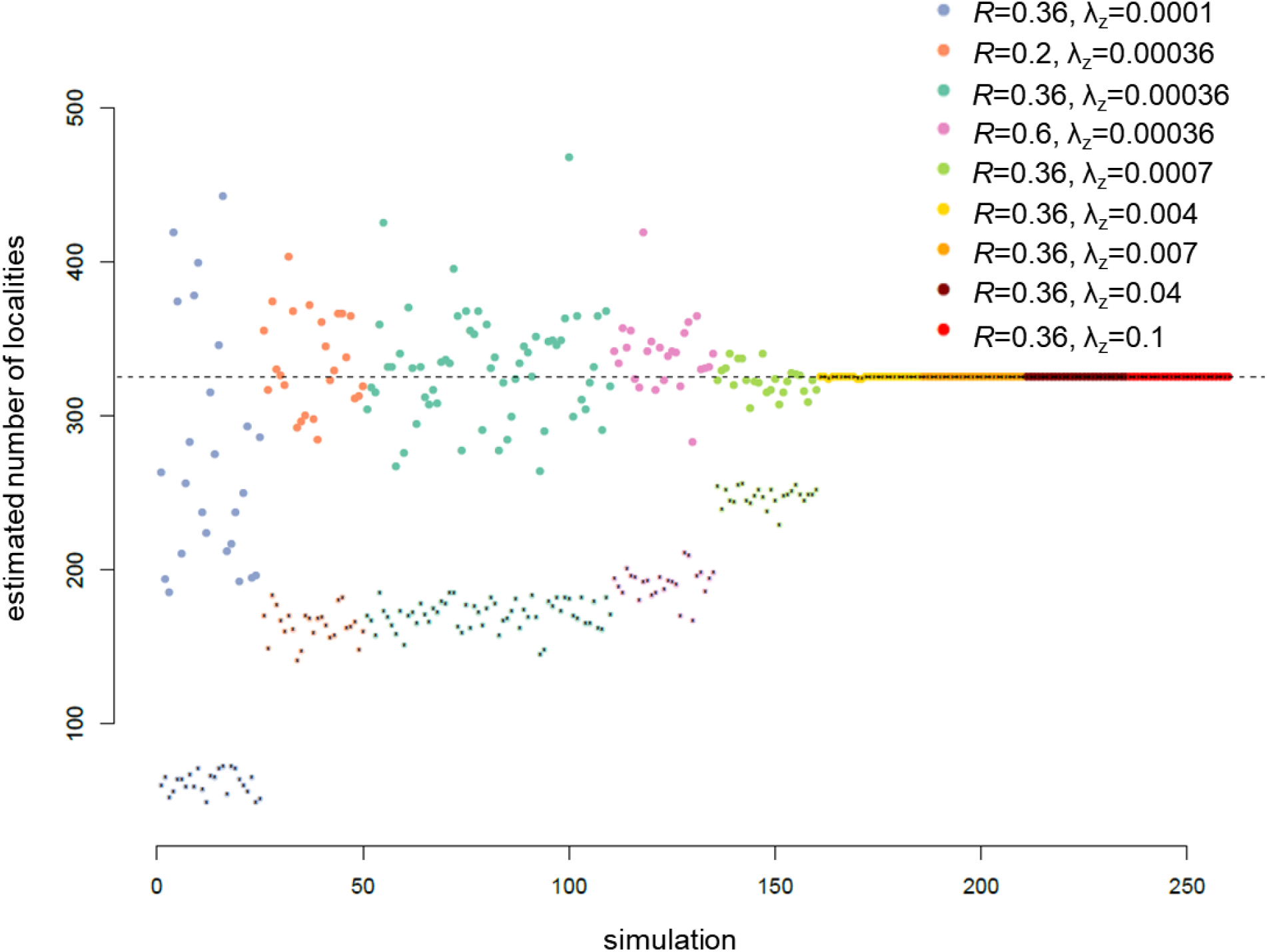
Estimated number of localities under surveillance. (calculated given the number of observed localities and the estimated parameter values). Large colored dots indicate the estimated number of localities under surveillance for each simulated dataset while the smaller dots show the number of localities observed in the dataset. The true number of localities is represented by the horizontal dashed line. Each color corresponds to a different parameter set used for simulations.

**S3 Fig.**
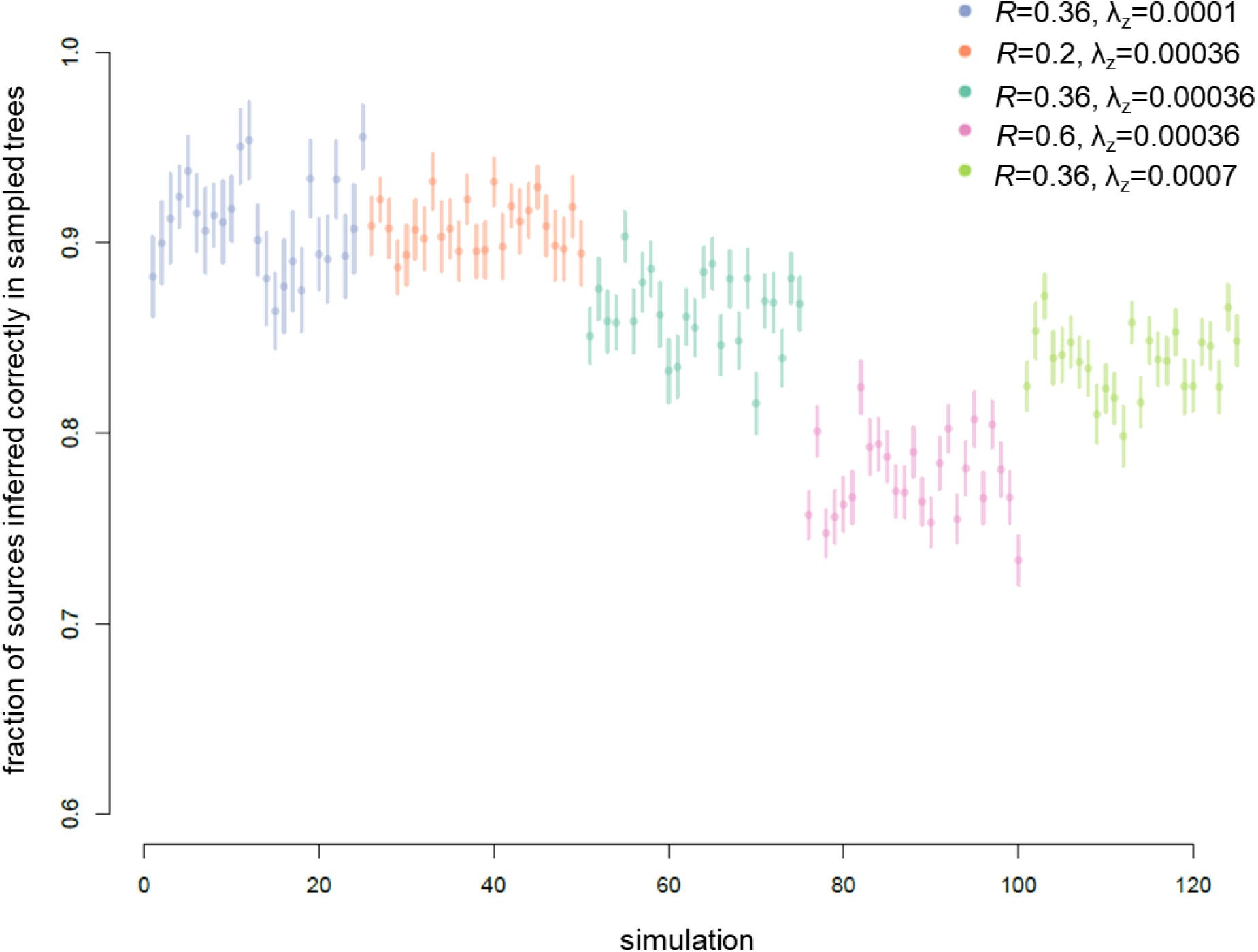
Accuracy of inferred transmission trees at inferring the correct source of cases. For each simulated dataset (25 simulations for each of 5 parameter sets), 200 transmission trees were drawn. Points show the mean fraction of cases inferred correctly in a sampled transmission tree and bars indicate the standard deviation.

**S4 Fig.**
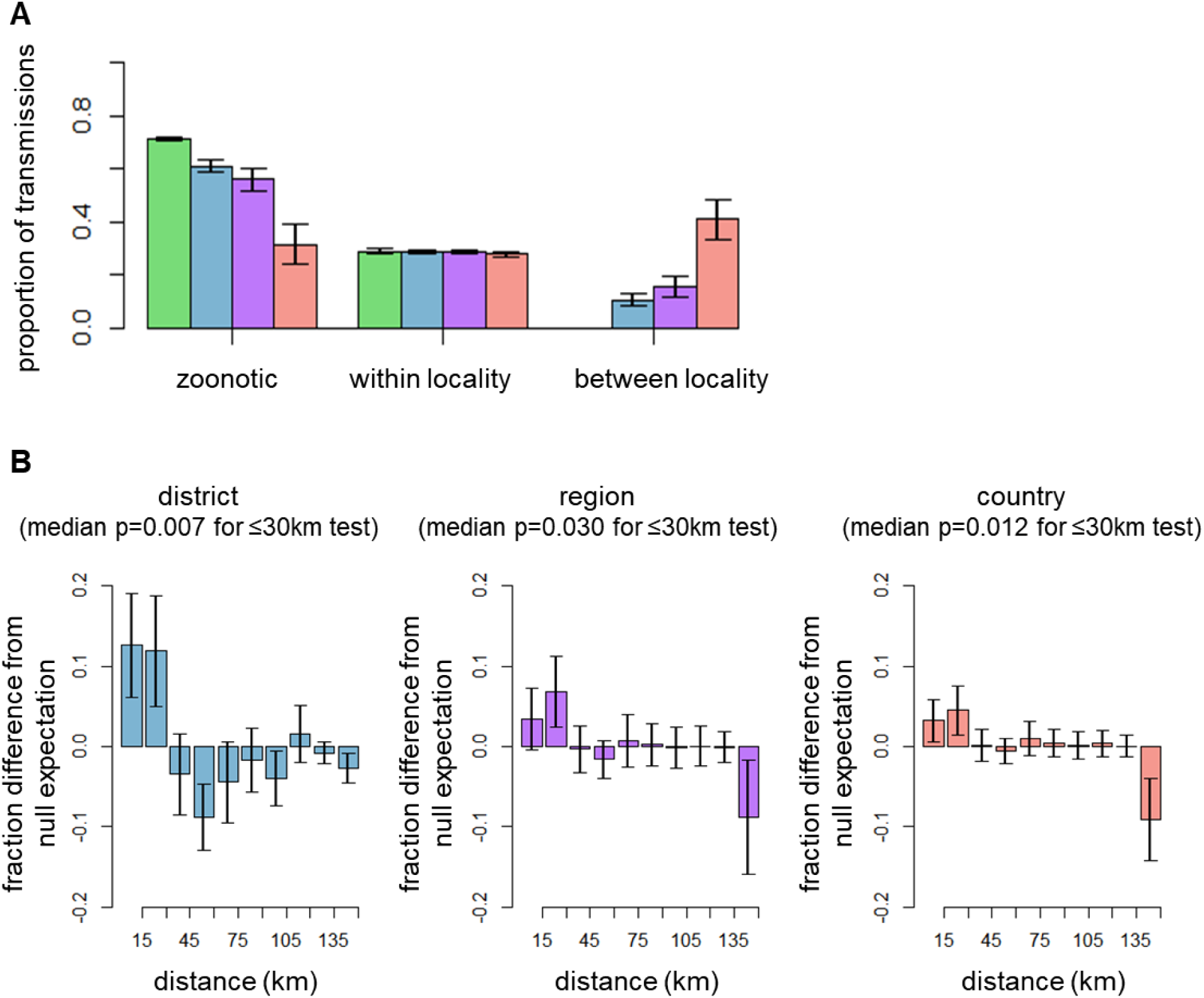
Inferred sources of monkeypox cases. **A.** The fraction of cases inferred to have originated from each source using each of the four spatial models (locality-green, district-blue, region-purple, country-red). **B.** Difference in the proportion of inter-locality human-to-human transmissions inferred by the models to occur over a given transmission distance versus expected based on the spatial distribution of localities. The p-values indicate the probability of observing as many or more transmissions over distances of ≤ 30 kilometers based on the null model (i.e. assuming distance plays no role in determining which localities are linked by inferred transmission events). The median p-value of sampled transmission trees is given, and the full distribution of p-values can be seen in S5 Fig.

**S5 Fig.**
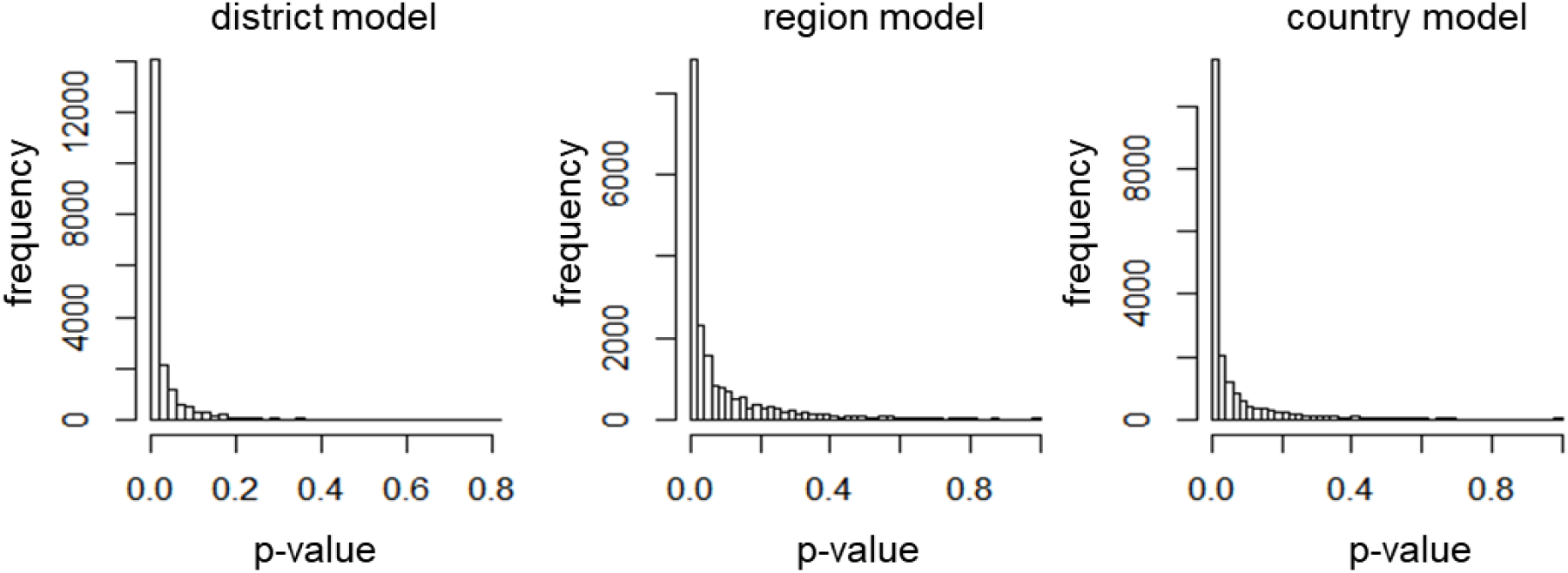
The distribution of p-values obtained across sampled transmission trees. P-values obtained from a binomial test examining whether the number of transmission events inferred to occur across thirty or fewer kilometers is greater than that expected based on the null distribution. Each p-value corresponds to a sampled transmission tree.

**S6 Fig.**
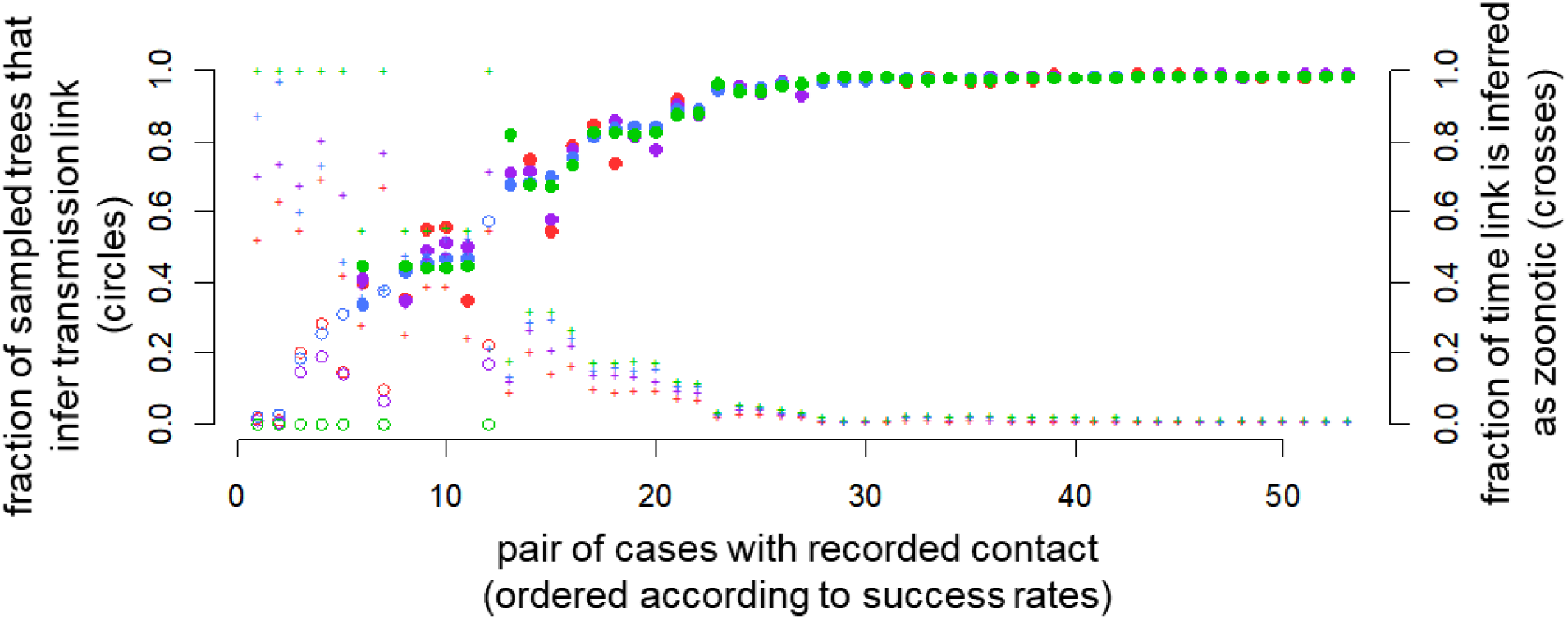
Comparison of epidemiologically contact-traced links with sampled transmission trees. C<u>ircles</u> (left axis) show the fraction of sampled trees that infer the epidemiologically-traced source. Open circles represent inter-locality links while closed circles represent intra-locality links. <u>Bars</u> (right axis) indicate the probability that a link is instead inferred to have a zoonotic source. Results are shown for models that use the country-level (red), region-level (purple), district-level (blue), and locality-level (green) broader contact zones. Links are sorted from lowest to highest success in the district model.

**S7 Fig.**
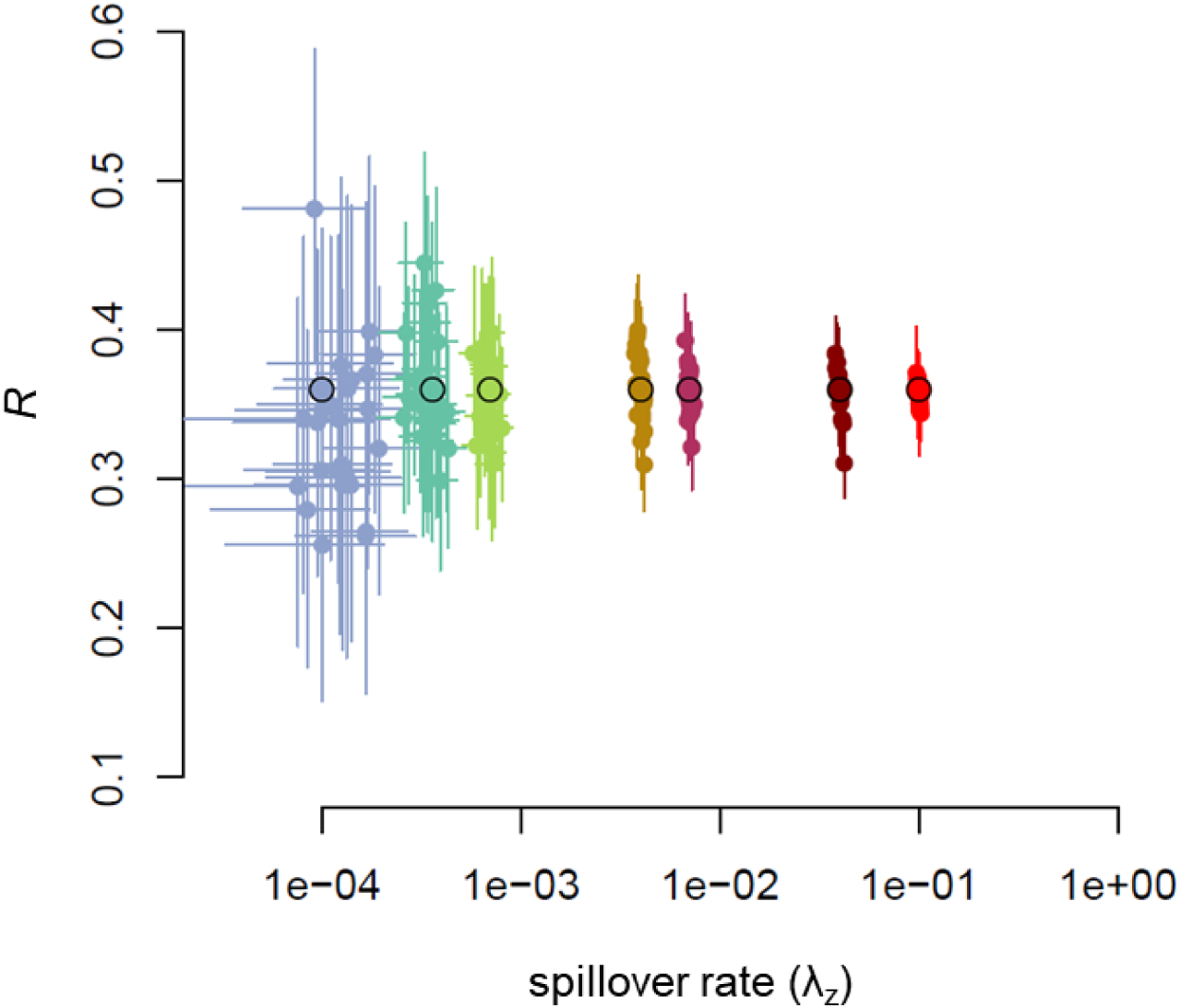
Effect of increasing spillover rate on parameter estimate success. Within each color, large points outlined in black indicate the true parameter set and smaller points indicate the inferred parameter values from 25 simulated datasets (lines show the 95% credible interval). Warmer colors correspond with higher spillover rates. Note the log-scale x-axis.

**S8 Fig.**
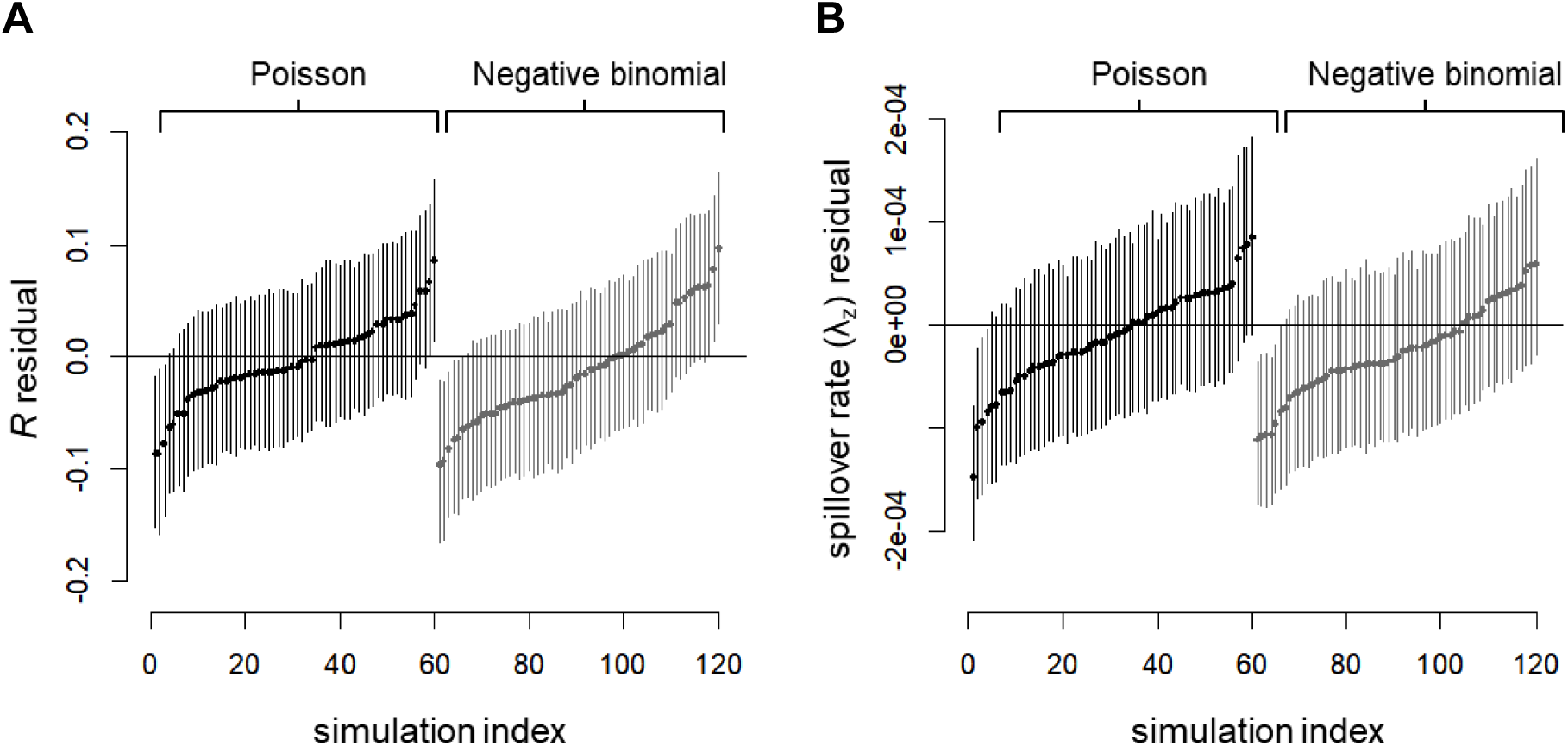
Parameter estimate residuals for data simulated using a negative binomial versus Poisson offspring distribution. Because the inference method assumes a Poisson offspring distribution, we compared the inference successes for datasets simulated assuming a Poisson offspring distribution versus datasets simulated assuming a negative binomial offspring distribution. The residuals in parameter estimates for 25 simulations are shown for A) the reproductive number and B) the spillover rate.

**S9 Fig.**
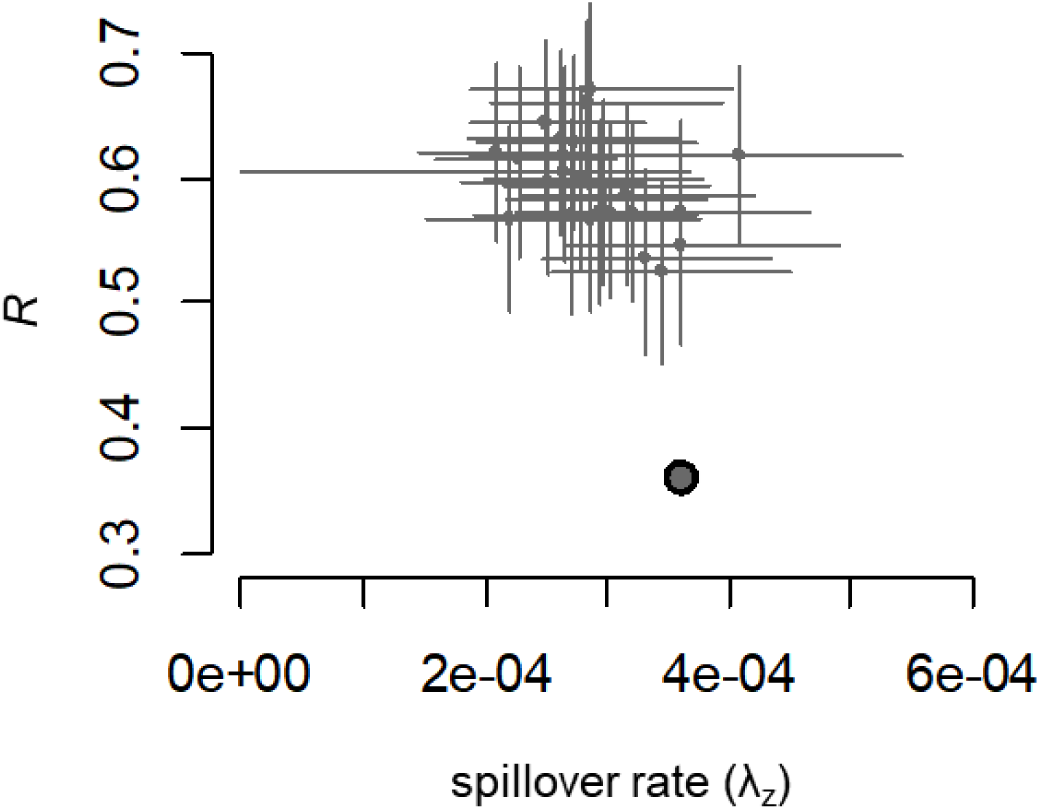
Strongly heterogeneous spillover causes bias in parameter estimates. The true parameter value is indicated by the large dot while smaller points indicate the inferred values from 25 simulated datasets (lines show the 95% credible interval). Simulations were conducted to mimic pockets of zoonotic disease moving through the reservoir population. To capture the idea that, at any given time, only a small subset of localities might be experiencing high levels of spillover while the rest of the localities experienced no spillover, the simulations assumed that every 25 days a new set of three localities experienced the full force of spillover for the entire system. This gave rise to clusters of primary cases, which tend to be misclassified as human-to-human transmission events by our inference approach, which assumes homogeneous spillover rates.

**S10 Fig.**
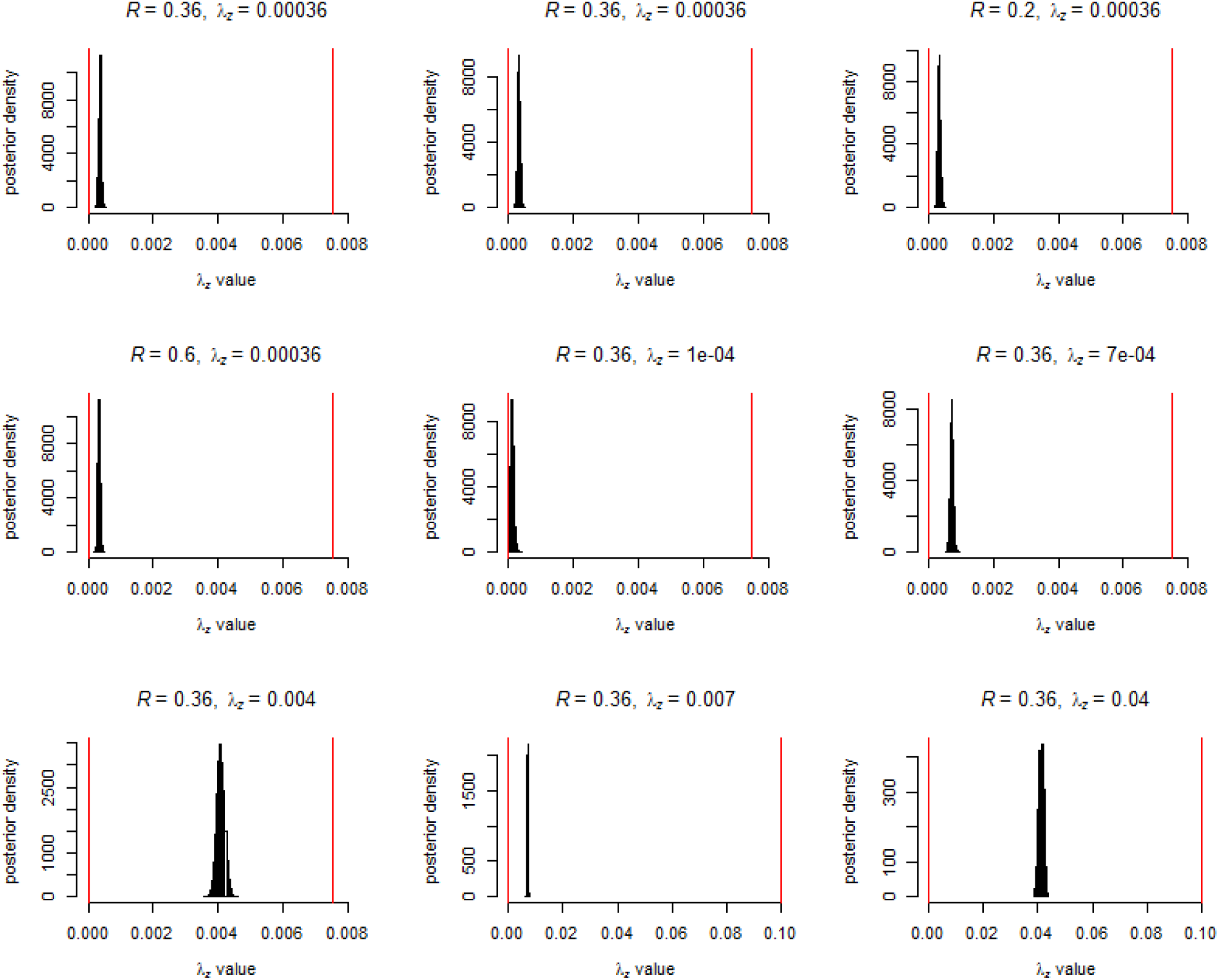
Comparison of prior and posterior distributions for spillover rate λ_z_. Black bars represent posterior distribution while red lines mark limits of the uniform prior distribution. One representative simulation is shown for each of the nine parameter sets. Notice that the posterior distribution is always relatively far from upper bound of the prior.

**S1 Table.**
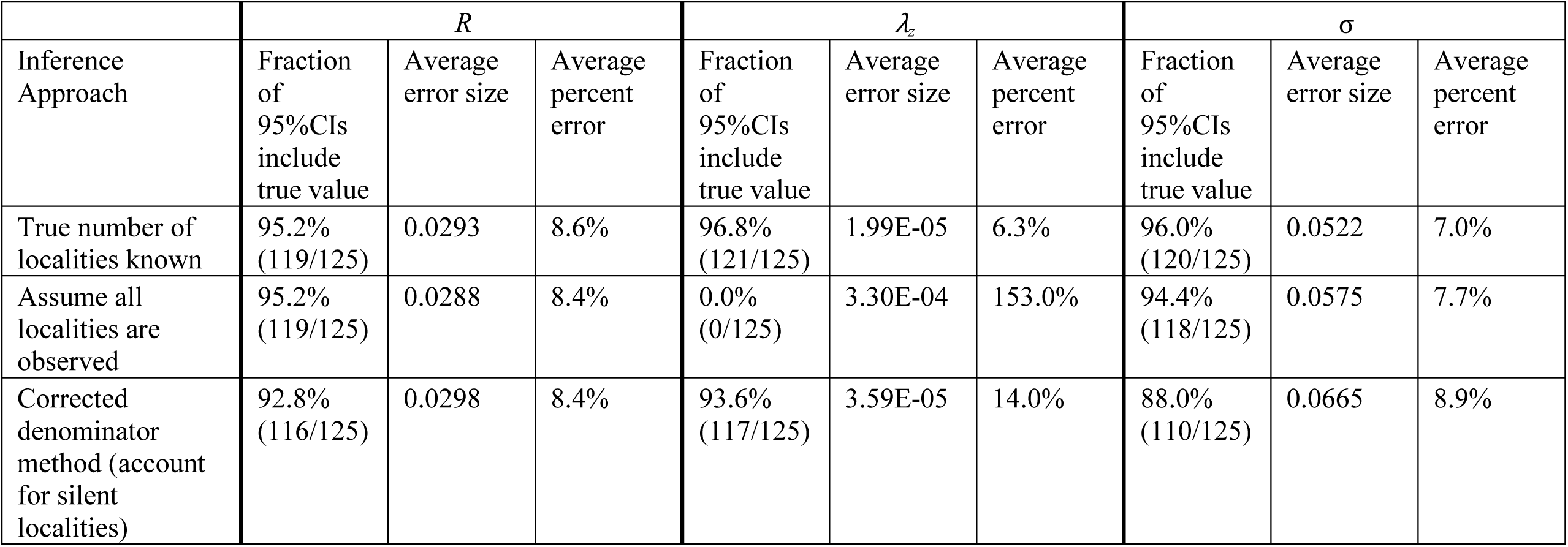
Comparison of inference method success over the same simulated datasets.

| Inference Approach | $R$ | | | $\lambda_z$ | | | $\sigma$ | | |
| --- | --- | --- | --- | --- | --- | --- | --- | --- | --- |
|  | Fraction of 95% CIs include true value | Average error size | Average percent error | Fraction of 95% CIs include true value | Average error size | Average percent error | Fraction of 95% CIs include true value | Average error size | Average percent error |
| True number of localities known | 95.2% (119/125) | 0.0293 | 8.6% | 96.8% (121/125) | 1.99E-05 | 6.3% | 96.0% (120/125) | 0.0522 | 7.0% |
| Assume all localities are observed | 95.2% (119/125) | 0.0288 | 8.4% | 0.0% (0/125) | 3.30E-04 | 153.0% | 94.4% (118/125) | 0.0575 | 7.7% |
| Corrected denominator method (account for silent localities) | 92.8% (116/125) | 0.0298 | 8.4% | 93.6% (117/125) | 3.59E-05 | 14.0% | 88.0% (110/125) | 0.0665 | 8.9% |

**S2 Table.**
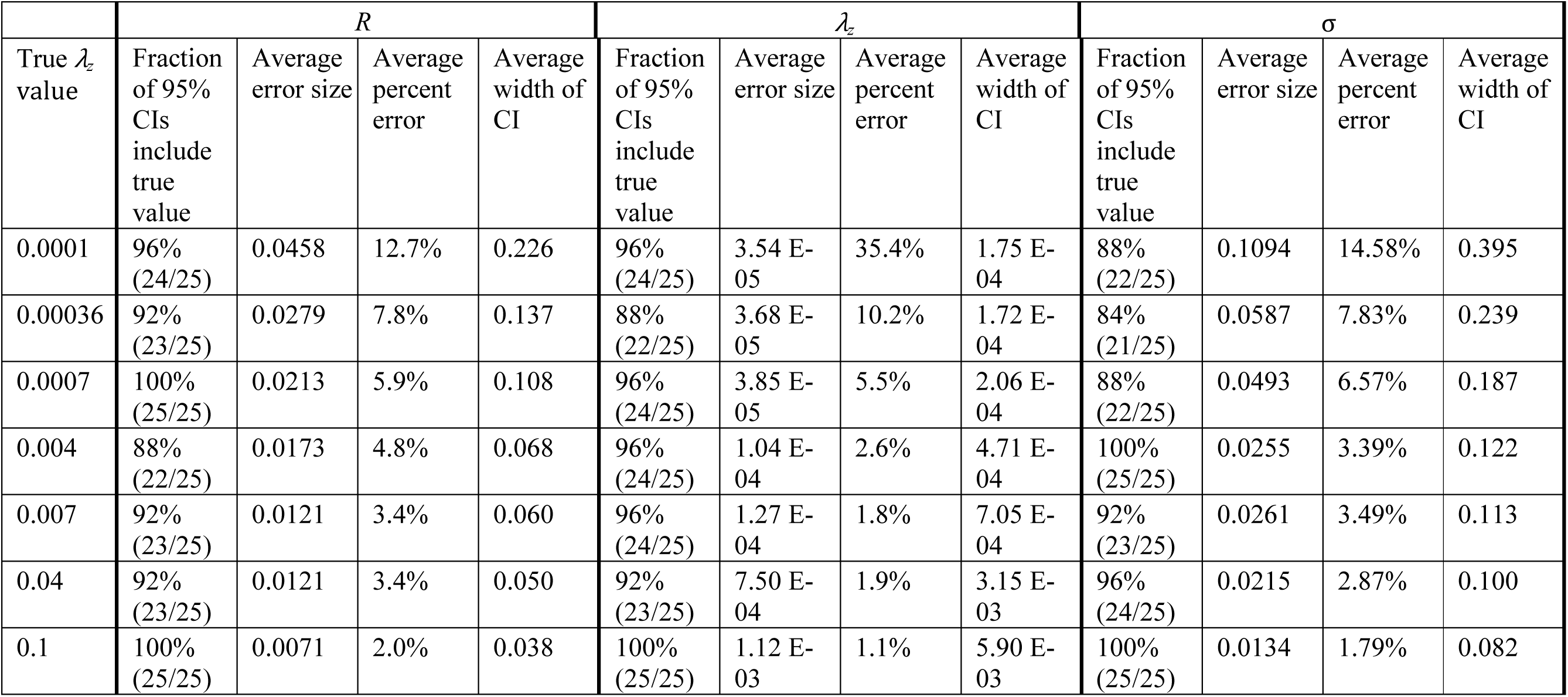
Success of the corrected denominator inference method for datasets simulated with increasing spillover rates.

**S3 Table.** Success of the corrected denominator inference method for datasets simulated with different offspring distributions.

| | $R$ | | | $\lambda_z$ | | | $\sigma$ | | |
| --- | --- | --- | --- | --- | --- | --- | --- | --- | --- |
| Offspring distribution | Fraction of 95% CIs include true value | Average error size | Average percent error | Fraction of 95% CIs include true value | Average error size | Average percent error | Fraction of 95% CIs include true value | Average error size | Average percent error |
| Poisson | 91.7%<br>(55/60) | 0.0289 | 8.0% | 93.3%<br>(56/60) | 3.76E-05 | 10.4% | 83.3%<br>(50/60) | 0.0649 | 8.7% |
| Negative binomial<br>($k=0.58$) | 86.7%<br>(52/60) | 0.0393 | 10.9% | 90.0%<br>(54/60) | 4.18E-05 | 11.6% | 91.7%<br>(55/60) | 0.0555 | 7.4% |

**S4 Table.** **Comparison of parameter estimates inferred using models of increasing spatial scale – data simulated using the ‘nearest five neighbors’ inter-locality transmission rule** where localities take the same GPS coordinates as in the DRC monkeypox surveillance dataset (true R is 0.36, true spillover rate is 0.00036; mean parameter estimates from inference on 25 simulated datasets).

| Model used for inference | mean $R$ | mean $\lambda_z$ |
| --- | --- | --- |
| District | 0.314 | 0.000346 |
| Region | 0.323 | 0.000343 |
| Country | 0.354 | 0.000328 |

**S5 Table.**
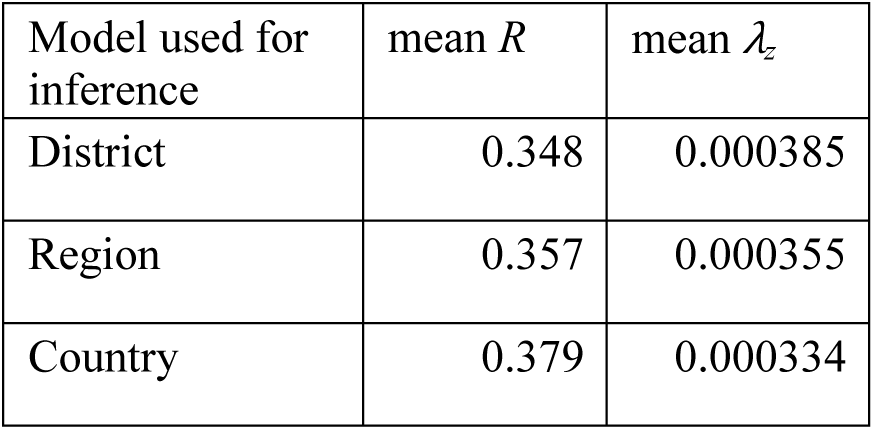
**Comparison of parameter estimates inferred using models of increasing spatial scale – data simulated assuming inter-locality transmission can occur between any localities located within 30 km of one another,** where localities take the same GPS coordinates as in the DRC monkeypox surveillance dataset (true R is 0.36, true spillover rate is 0.00036; mean parameter estimates from inference on 25 simulated datasets).

| Model used for inference | mean $R$ | mean $\lambda_z$ |
| --- | --- | --- |
| District | 0.348 | 0.000385 |
| Region | 0.357 | 0.000355 |
| Country | 0.379 | 0.000334 |

**S6 Table.**
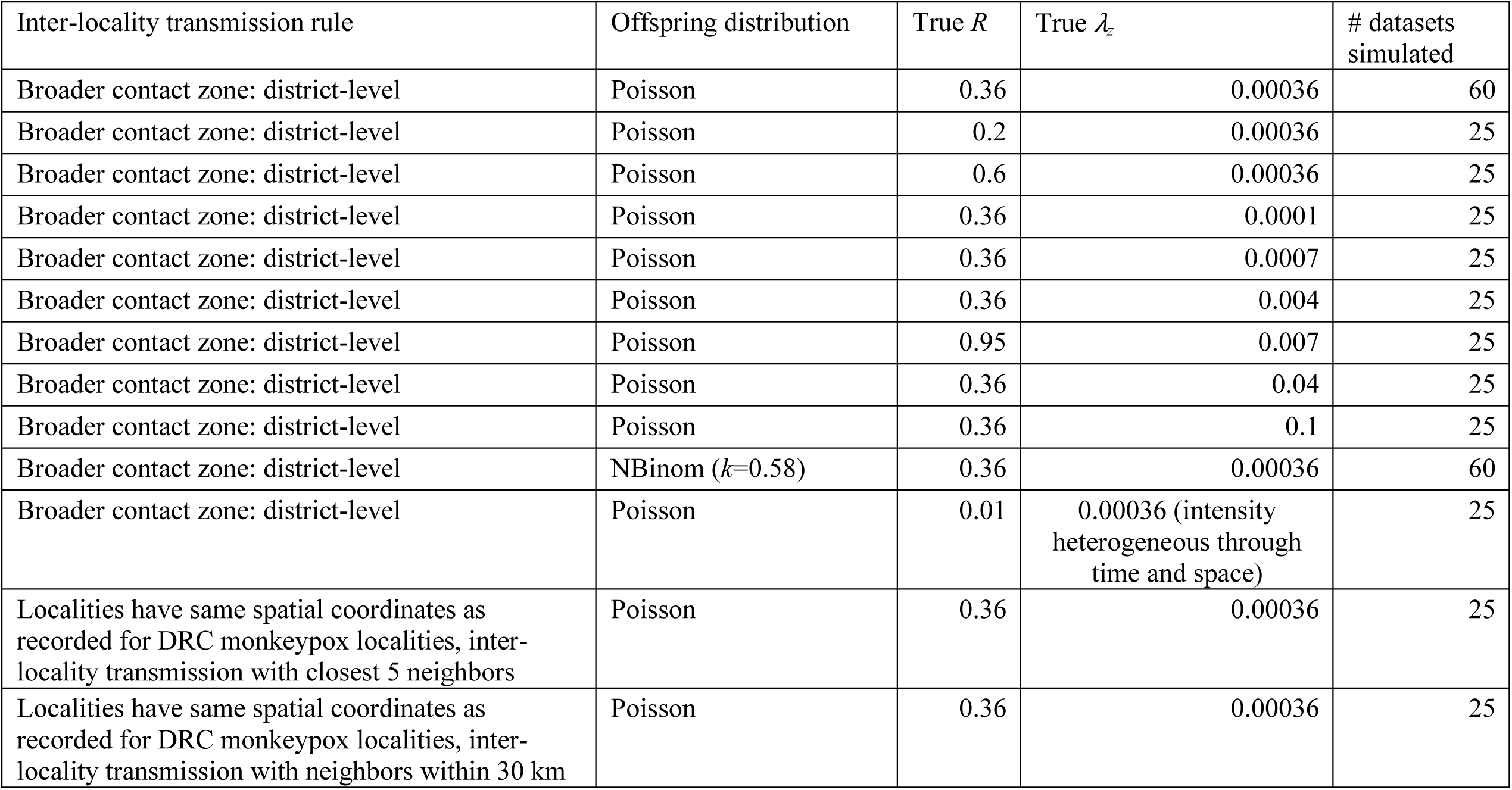
Description of datasets simulated.

**S7 Table.** Parameter descriptions.

| Symbol | Description |
| --- | --- |
| $\mu_{t,v}$ | Expected number of cases observed on day $t$, in locality $v$ |
| $N_{t,v}$ | Actual number of cases observed on day $t$, in locality $v$ |
| $N$ | Actual number of cases observed across all localities over the course of surveillance |
| $V$ | Total number of localities under surveillance |
| $V_w$ | Total number of localities under surveillance in the broader contact zone of locality $w$ |
| $W$ | Number of localities with one or more cases (the number of localities that appear in the surveillance dataset) |
| $W_w$ | Number of localities with one or more cases in the broader contact zone of locality $w$ |
| $T$ | Duration of surveillance: number of days surveillance was conducted |
| $\lambda_z$ | Spillover rate: the expected number of spillover events per day in a given locality |
| $\lambda_{\{s,w\},\{t,v\}}$ | The expected number of new infections that become symptomatic on day $t$ in locality $v$ caused by an infectious individual who became symptomatic on day $s$ in locality $w$ |
| $R$ | Reproductive number: the average number of secondary cases caused by an infectious individual |
| $\sigma$ | Within-locality transmission proportion: the fraction of cases arising from human-to-human transmission that occur in the same locality as the source case |

**S1 Data. Case records.** For all individuals included in the analyses, records the case identification number, the locality identification number, the day of surveillance when disease onset occurred (the first day of fever when known, otherwise the first day of the rash), the names of the district and region where the case occurred, and masked GPS coordinates of the locality. The geographic masking technique known as ‘donut masking’ was used to obscure the exact location of cases and preserve privacy. For each locality with a recorded location, two random values were drawn: the first determines the direction and the second determines the distance from the original point. The new location is within 0.1 degrees from the original point but not closer than 0.02 degrees.

**S2 Data. Contact-tracing links.** Each row provides the case identification numbers for a pair of cases that was identified as a probable transmission link through epidemiological contact-tracing.

## S1 Text. Supplementary material on methods

### Corrected denominator method: Derivation for the conditional likelihood function

The model described in the main text tells us that the number of new human cases on day *t* in locality *v* follows a Poisson distribution with mean

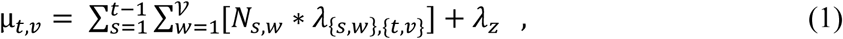

which represents the sum of the expected numbers of cases caused by spillover and all previous human cases (S7 Table provides a description of parameters). Based on this model, the likelihood of a set of parameters (*θ* = {*R*, *λ_z_, σ*}) given surveillance data (*D* = *N_t,𝒱_* cases observed on each day *t* and locality *v*) is:

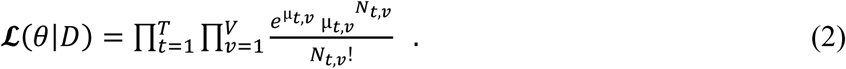

A challenge in applying this likelihood function to surveillance data arises when the total number of localities under surveillance, *V*, is unknown. Instead, we observe *W* localities that have one or more observed cases. If we re-arrange the product functions in the likelihood function, it becomes more apparent that we are taking the product of the likelihood for each locality:

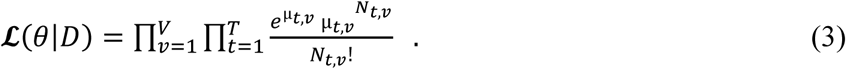

However, because we only observe localities with one or more cases in the surveillance data, we need that conditioning to be reflected in the likelihood. In other words, we now want to express the likelihood of a particular time-series of cases in a locality *conditional on that locality having one or more cases*. This can be done for each locality by multiplying its component of the likelihood by the inverse of the probability (*q*) of having one or more cases:

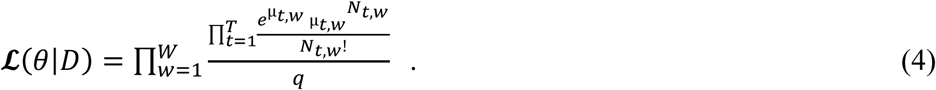

It is now necessary to calculate the probability a surveilled locality experiences one or more cases. This probability is equivalent to one minus the probability of no cases occurring at a locality during the surveillance period. The following section explains how the probability of zero cases occurring at a given locality (here denoted *p*) is calculated.

For zero cases to occur in a locality, there must be no zoonotic spillover into that locality as well as no human-to-human transmission from an outside locality. The zoonotic component is relatively straightforward to calculate, as it is simply the probability of zero spillover events on each of the *T* days (which equals *e*^−*λ_z_T*^). The probability of no transmission from an outside human source is a bit more complicated and can be broken down by the generation of the outside case to avoid double-counting. The generation of a case indicates how many human-to-human transmission events occurred leading to the case. We refer to cases resulting from zoonotic spillover as primary cases. Individuals infected by primary cases are second generation cases, individuals infected by second generation cases are third generation cases, etc. For there to be no cases in a locality, no transmission may have occurred into that locality from outside cases in any generation:

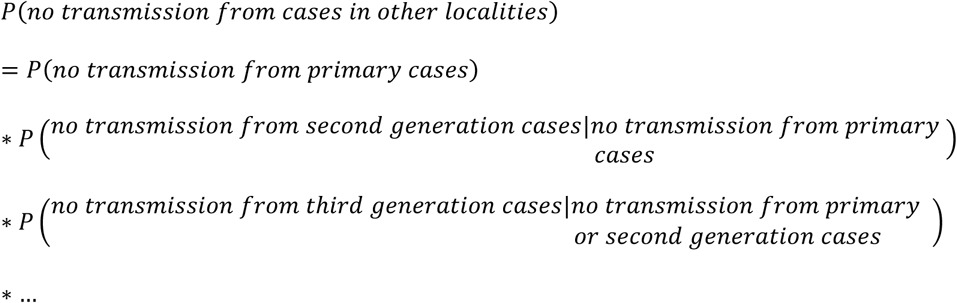

The number of cases caused by a given case (of any generation) in the target locality is described by a Poisson distribution with expected value equal to 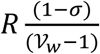, where *V_w_* is the number of localities within the target locality’s broader contact zone. Because each case transmits disease independently of one another (conditioned on the previous cases), the probability that no generation *i* cases cause infections in the target locality is 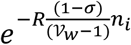, where *n_i_* is the total number of *i*^th^ generation cases within the broader contact zone (given knowledge that none of the cases from previous generations transmitted to the target locality). Incorporating this information, the probability of observing zero cases in a locality (*p*) becomes:

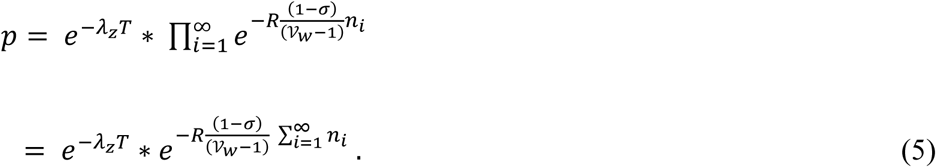

We next need to calculate estimates for the expected values of each of the *n_i_*. The expected number of primary cases in the entire broader contact zone (given that no spillover events occurred into the target locality) is the expected number of spillover events per locality (*λ_z_*) multiplied by the number of localities under consideration (𝒱*_w_* − 1), multiplied by the number of surveillance days (*T*). For subsequent case generations, we can calculate the expected number of cases in generation *i*+1 as the number of cases caused by the *i*^th^ generation in their own localities plus those caused in the 𝒱_*w*_ − 2 other possible localities (there are 𝒱*_w_* − 2 other possible localities because the case’s current locality and the target locality have already been counted):

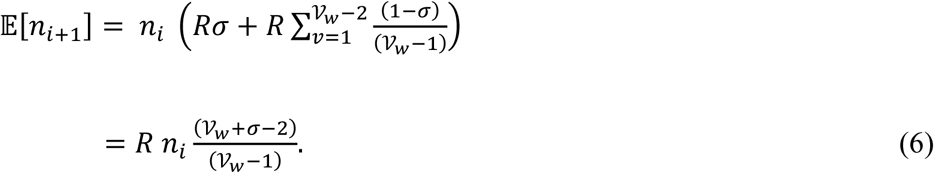

If we approximate the values of *n_i_* with 𝔼[*n_i_*], we get

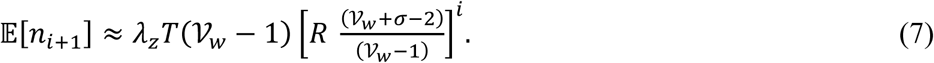

Returning to our estimation of *p*, we can approximate *n_i_* values with 𝔼[*n_i_*] and get

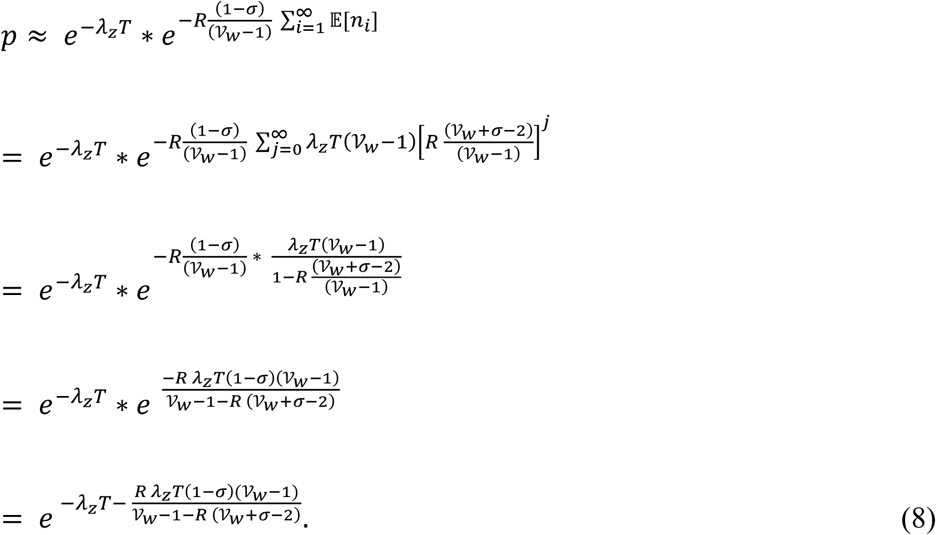

With some additional algebraic simplification, we can insert this value in the original equation:

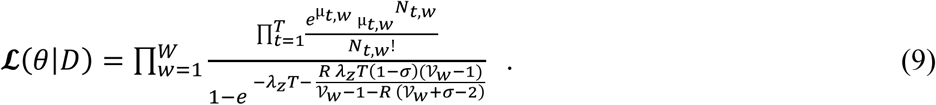

This expression still includes the parameter 𝒱_*w*_, though fortunately the sensitivity of results to the value of this parameter is relatively low. We therefore approximate 𝒱_*w*_ using the expected number of localities under surveillance in the broader contact zone. This calculation is explained in the following section.

### Estimating total number of localities under surveillance

We wish to use the estimated parameter values for *R*, *λ_z_*, and *σ* in conjunction with the number of observed localities in a broader contact zone (*W_w_*) to estimate the total number of localities under surveillance in that broader contact zone (*V_w_*). If we let *q* be the probability a locality is observed (has one or more cases during the surveillance period), then we expect *V_w_* \**q* ≈ *W_w_*. From the section above, we approximate *q* = 1-*p* as:

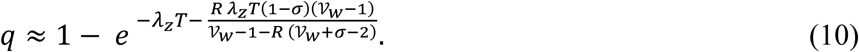

So we estimate *V_w_* as the value that satisfies the equation:

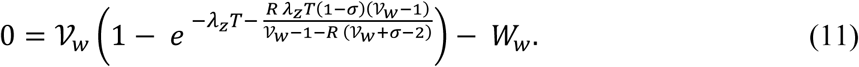

### Simulation methods

#### Simulations with exact spatial locations

Although the model assumes that inter-locality transmission with a broader contact zone is equal between all locality pairs, we expect that the actual amount of shared transmission between two localities is strongly influenced by the distance between those localities. We conducted two simulations using localities with set geographic locations and inter-locality transmissions depending on the spatial relationship of the localities. We took the 178 GPS records available from monkeypox surveillance in the DRC during the 1980s and simulated transmission across localities with the same coordinates and the same district and region boundaries. Two types of inter-locality transmission rules were explored. In the first of these, inter-locality transmissions were assumed to occur equally into a source locality’s five closest neighbors. In the second set of simulations, inter-locality transmissions from a source locality were assumed to occur equally among all outside localities within 30 km of the source locality.

#### Simulations with highly structured and non-homogeneous spillover patterns

To illustrate how highly structured and non-homogeneous spillover could bias parameter estimates, we simulated an extreme case of a zoonotic epidemic traveling through time and space. We imagined that disease dynamics in the reservoir would occur in a single location for 25 days before moving to a new spot, in an extreme form of a traveling zoonotic epidemic. For each 25 day period, three localities (selected to be in the same district when possible) would be selected to experience all of the spillover in the entire system. Aside from this extreme spillover pattern, the simulation followed the district-level model.

### Sensitivity analyses

#### Sensitivity of parameter inference to elevated or heterogeneous spillover

To test whether a high rate of spillover would inundate the system with so many cases that the temporal clustering patterns resulting from human-to-human transmission could be obscured, we simulated datasets with spillover rates up to 0.1. This value corresponds with an expected 59,312.5 spillover events during the five year simulation, which corresponds to an average of 36.5 per year in each locality. At this rate of spillover, there is an average of only ten days between spillover events, a shorter period than the mean generation time for human-to-human transmission events, which was sixteen days. Across the range of spillover rates tested, the method did very well at both point estimates and capturing the true parameter values within the 95% CI (an average of 94.3% of CIs included the true value of *R* and 94.9% included the true value of *λ_z_;* S7 Fig, S2 Table). As the spillover rate increased from 0.0001 to 0.1, estimates of *R* tended to improve (posterior means closer to true value and smaller CIs). While the absolute error on estimates of *λ_z_* increased as spillover rate increased, the relative error tended to decrease. As such, it appears that elevated spillover rates, far from obscuring patterns, may actually correspond with improved estimates, presumably due to the increased inference power resulting from a larger number of cases.

Spillover is unlikely to occur homogeneously through time and space in real-world settings. As an illustration of the potential effect this occurrence could have on parameter estimates, we simulated an extreme case (see ‘Simulations with highly structured and non-homogeneous spillover patterns,’ above) where spillover occurs into three localities at a time. The parameter inference results for this situation were strongly biased (S10 Fig).

#### Sensitivity of parameter inference to offspring distribution assumptions

The model used in this study assumes that the number of new cases caused by an infectious individual follows a Poisson distribution, but previous work suggests that the offspring distribution is often better characterized by a negative binomial distribution, which allows for a greater amount of variation between individuals [1]. We simulated datasets using a negative binomial offspring distribution (using a dispersion parameter *k*=0.58 in accordance with previous estimates for monkeypox from [1]) and examined how well our inference method, which assumes a Poisson offspring distribution, estimated the true parameter values. Estimates for these datasets were only marginally less accurate than estimates for datasets generated with a Poisson offspring distribution (with an average percent error of 10.9% as opposed to 8.2% for *R* and of 11.6% as opposed to 10.4% for spillover rate estimates) (S8 Fig, S3 Table). As such, there are unlikely to be strong biases introduced from a mis-specified offspring distribution for the monkeypox dataset, though this bias could increase if applied to pathogens with more extreme transmission variance.

#### Sensitivity of parameter inference to broader contact zone assumption

To examine how assuming different broader contact zones would affect inference results, we compared parameter estimates obtained under three choices of broader contact zones for data simulated under two inter-locality transmission rules. We simulated disease spread in a system where localities were placed in the same arrangement as seen in 178 localities with GPS coordinates included in the monkeypox surveillance system, district and region arrangement were the same as in the 1980s surveillance, and human-to-human transmission could occur either between a locality and its five closest neighbors or between localities located within 30 km of one another. Inference results again showed increasing estimates of *R* and decreasing estimates of spillover rate as the size of the assumed broader contact zone increased (S4 and S5 Table).

